# Automated liquid handling extraction and rapid quantification of underivatized amino acids and tryptophan metabolites from human serum and plasma using dual-column U(H)PLC-MRM-MS and its application to prostate cancer study

**DOI:** 10.1101/2023.12.31.573763

**Authors:** Tobias Kipura, Madlen Hotze, Alexa Hofer, Anna-Sophia Egger, Lea E. Timpen, Christiane Opitz, Paul A. Townsend, Lee A. Gethings, Kathrin Thedieck, Marcel Kwiatkowski

## Abstract

Free amino acids (AAs) and their metabolites are important building blocks, energy sources and signaling molecules associated with various pathological phenotypes. The quantification of AA and tryptophan (TRP) metabolites in human serum and plasma is therefore of great diagnostic interest. Robust and reproducible sample extraction and processing workflows as well as rapid, sensitive absolute quantification of AA and TRP metabolites are required to identify candidate biomarkers and to improve current screening methods. We developed a validated semi-automated extraction and sample processing workflow using a robotic liquid handling platform and a rapid method for the absolute quantification of 20 free, underivatized AAs and 6 TRP metabolites using dual-column U(H)PLC-MRM-MS. The automated extraction and sample preparation workflow is designed for use in a 96-well plate format, allowing robust and reproducible high sample throughput without the need for further SPE, evaporation and/or buffer exchange. Samples extracted from serum and/or plasma in 96-well plates can be transferred directly to the U(H)PLC autosampler. The dual-column U(H)PLC-MRM-MS method, using a mixed-mode reversed-phase anion exchange column with formic acid as mobile phase modifier and a high-strength silica reversed-phase column with difluoroacetic acid as mobile phase additive, provided absolute quantification with nanomolar lower limits of quantification (LLOQ) for all metabolites except glycine (LLOQ: 2.46 µM) in only 7.9 minutes. The semi-automated extraction workflow and dual-column U(H)PLC-MRM-MS method was applied to a human prostate cancer study and was shown to discriminate between treatment regimens and to identify amino acids responsible for the statistical separation between healthy controls and prostate cancer patients on active surveillance.

## Introduction

Amino acids (AAs) are the building blocks of proteins and other metabolites, including nucleotides and hormones, and serve as energy sources and signaling molecules. In addition, metabolites derived from AAs, such as those derived from tryptophan metabolism (TRP metabolites), play important roles in the regulation of inflammation, neurological functions, immune responses and tumor metabolism ^1–3^. Alterations in AA levels are known to result from defects in AA transport ^4^, AA metabolism ^5^ or dietary conditions ^6^, and are associated with pathological phenotypes ^7–11^. The monitoring AA and its metabolites in human blood-derived samples such as serum and plasma is therefore of great diagnostic and clinical interest and can contribute to the identification and validation of new biomarker candidates and/or improve current screening methods.

The identification of candidate biomarkers and their validation generally requires large sample sizes. Sample extraction is one of the most critical steps in such studies and experiments, as it is usually the first experimental step on which the outcome of the experiment largely depends. Manual sample extraction of large sample quantities is a challenging, time-consuming and psychologically stressful task for laboratory personnel, as the outcome of the experiment can be greatly affected by this step. The quality of sample collection and preparation can be highly dependent on the individual and/or daily performance of the operator and, especially when multiple operators are involved, can affect the quality and comparability of the data. Automated and/or semi-automated sample extraction and sample preparation workflows can mitigate these issues. Compared to manual pipetting, automated liquid handling can increase robustness, reproducibility, accuracy, traceability, optimized chemical consumption and throughput ^12^. In recent years, miniaturized liquid handling platforms and robotic pipetting systems have been developed that are highly modular, flexible and easily programmable. This provides a simpler and more intuitive approach to developing laboratory workflows that enable automated, robust, reproducible and traceable sample processing of large sample volumes. As this is a relatively new development, there have been few publications in the life science community describing the development and validation of automated sample preparation with liquid handling platforms and making them available to the community ^13,14^.

In addition to reproducible sample extraction, simple, rapid, sensitive and selective methods are required for the quantification of amino acids and TRP metabolites in complex biological matrices such as human serum and plasma. However, the wide variety of structures and physicochemical properties of these metabolites make them challenging to analyze. TRP metabolites are usually analyzed by liquid chromatography (LC) or gas chromatography (GC) using various detection methods such as UV absorption, fluorescence and electrochemical methods ^15^. However, LC and GC methods using UV/VIS, fluorescence and electrochemical detectors are not suitable for comprehensive or universal detection of TRP metabolites at their physiological concentrations in biological samples. For this reason, LC methods with mass spectrometric detection (LC-MS) have been increasingly developed in recent years ^16–18^, as mass spectrometers are the closest to a “universal detector” for the analysis of TRP metabolites at their physiological concentrations. AAs are small, polar, zwitterionic metabolites that have low ultraviolet (UV) absorption and most lack chromophores ^19^. AAs have therefore traditionally been analyzed by amino acid analyzers using pre- and/or post-column derivatization to enhance chromatographic separation and photometric detection by UV/VIS and fluorescence ^20,21^. The current gold standard for AA analysis, ion exchange (IEX) chromatography combined with post-column ninhydrin derivatization and fluorescence detection, is typically costly and time-consuming with run times of 1-3 hours, reducing sample throughput ^22,23^. LC-MS provides rapid, sensitive and selective analysis of amino acids. However, reversed-phase (RP) LC-MS applications are often problematic for AA analysis due to the low retention of AAs caused by their hydrophilicity and polarities. Therefore, many LC-MS approaches use pre-column derivatization, such as 6-aminoquinolyl-N-hydroxysuccinimidyl carbamate (AQC) ^24^ or chloroformate derivatization ^25^, to improve retention and separation. However, when analyzing AAs in complex biological samples, derivatization can increase interference due to derivative instabilities, reagent interferences, and side reactions with other matrix substances, thus reducing precision and accuracy while increasing errors, sample complexity, and sample preparation time. Non-derivative LC-MS approaches would therefore be advantageous. Ion pairing reagents, such as perfluorinated carboxylic acids (e.g., pentafluoropropionic acid, heptafluorobutyric acid, nonafluoropentanoic acid), can be used to improve AA retention in RP-LC-MS ^22,26,27^. Ion pairing reagents have the disadvantage of causing ionization suppression and retention time shifts, resulting in the LC-MS system being dedicated to a single polarity and type of analyte analysis, and long LC equilibration times ^28^. Hydrophilic interaction chromatography (HILIC) is an interesting alternative to RP-LC-MS and ion-pairing RP-LC-MS. However, HILIC often suffers from peak broadening, long LC equilibration times, limited robustness and insufficient separation of isobaric amino acids, such as isoleucine and leucine ^22,29,30^. The introduction of new RP and mixed-mode (MM) column chemistries specifically designed for the separation of polar analytes provide powerful alternatives to ion-pairing RP and HILIC for the analysis of TRP metabolites and free, underivatized amino acids by LC-MS^31–35^.

In this study, we present a semi-automated workflow for the analysis of underivatized free amino acids and TRP metabolites in human serum and plasma samples. AAs and TRP metabolites were extracted using a robotic liquid handling platform, minimizing manual intervention and maximizing reproducibility, and a dual-column ultrahigh performance liquid chromatography-multiple reaction monitoring mass spectrometry (U(H)PLC-MRM-MS) method enabled rapid and sensitive absolute quantification of 20 amino acids and 6 TRP metabolites in 7.9 min. The method was shown to discriminate between treatment regimens and identify amino acids responsible for the statistical separation of patient groups in a human prostate cancer study.

## Materials and Methods

### Materials

LC-MS grade methanol (MeOH), acetonitrile (ACN) and formic acid (FA) were purchased from Thermo Fisher Scientific (Dreieich, Germany). HPLC-grade trichloroacetic acid (TCA) and difluoroacetic acid (DFA) were purchased from Merck and Sigma-Aldrich (Munich, Germany). Water was purified using a Milli-Q purification system from Merck (Munich, Germany). Unlabeled metabolite standards were purchased from Sigma-Aldrich (Munich, Germany), stable-isotope labeled internal standards from Eurisotop (Saarbrücken, Germany, part of Cambridge Isotope Laboratories). Plasma and serum samples used for method development and method validation were obtained from the German Cancer Research Center (Deutsches Krebsforschungszentrum DKFZ, Heidelberg, Germany). The blood samples were obtained after informed consent and approval of the local regulatory authorities (Ethics Board approval S-496/2014). NIST SRM 1950 (www.nist.gov/srm) reference plasma was purchased from Sigma-Aldrich (Munich, Germany). Prostate cancer serum samples (University of Surrey, Guildford, UK) from 47 individuals were pooled into phenotypic groups. Sample pools were approved by the Yorkshire and the Humber-Leeds East Research Ethics Committee, UK, under reference number 08/H1306/115+5 and IRAS project ID 3582. Serum pools consisted of healthy controls (n = 8), a no-cancer group (individuals diagnosed with an infected prostate but without prostate cancer diagnosis) (n = 8), patients on active surveillance (n = 6), brachytherapy (n = 9), hormone therapy alone (n = 6), combined radiotherapy and hormone therapy (n = 4), and prostatectomy (n = 6), as previously described by Munjoma et al ^36^.

### Preparation of metabolite standards

The AAs were dissolved in 0.1 N HCl. 3-hydroxykynurenine (3OHKYN), kynurenine (KYN), nicotinic acid (NICAC) and nicotinamide (NICAM) were dissolved in 0.1% FA. Kynurenic acid (KYNAC) was dissolved in DMSO and indole-3-acetic acid (I3AA) was dissolved in 50% MeOH, 0.1% FA. The 20 AAs and 6 TRP metabolites were pooled and a dilution series was prepared from 6.25 µM to 0.05 µM dissolved in 0.1% FA and spiked with stable isotope labeled standards (ISTD) (Table S1) to achieve a concentration of 1.25 µM of ISTD in each sample of the dilution series. The dilution series was used during method development to determine linearity, lower limit of detection (LLOD), and lower limit of quantification (LLOQ), and as an external calibration series during patient sample analysis. For patient sample analysis, the added ISTDs in the calibration series were used to determine response ratios, which were used for absolute quantification of the analytes. To determine accuracy and precision, 5 QC samples dissolved in 0.1% FA were prepared at concentrations of 0.19 µM (QC LLOQ), 0.39 µM (QC low), 3.12 µM (QC medium), 4.69 µM (QC high) and 6.25 µM (QC ULOQ). QC and standard samples were spiked with ISTDs (Cambridge Isotope Laboratories, Table S1) to a final concentration of 1.25 µM to calculate the response ratio (peak area 12C/peak area 13C).

### U(H)PLC-MRM-MS

Free AAs and TRP metabolites (1 µL injection volume) were separated on an UPLC™ system (ACQUITY™ Premier, Waters, Milford, MA) using two columns in the column manager, namely an Atlantis™ Premier BEH™ C18 AX column (2.1x150 mm, 1.7 µm, Waters) maintained at 45°C and an ACQUITY UPLC HSS T3 column (optimized method: 2.1x50 mm, 1.8 µm; column screening: 2.1x150 mm, 1.8 µm, Waters) maintained at 35°C. Metabolites were separated using a H20:ACN gradient with 0.1% FA or 0.05% DFA as mobile phase modifier. The gradients used in the initial column screening and in the optimized method are shown in Table 1 below.

**Table 1:**
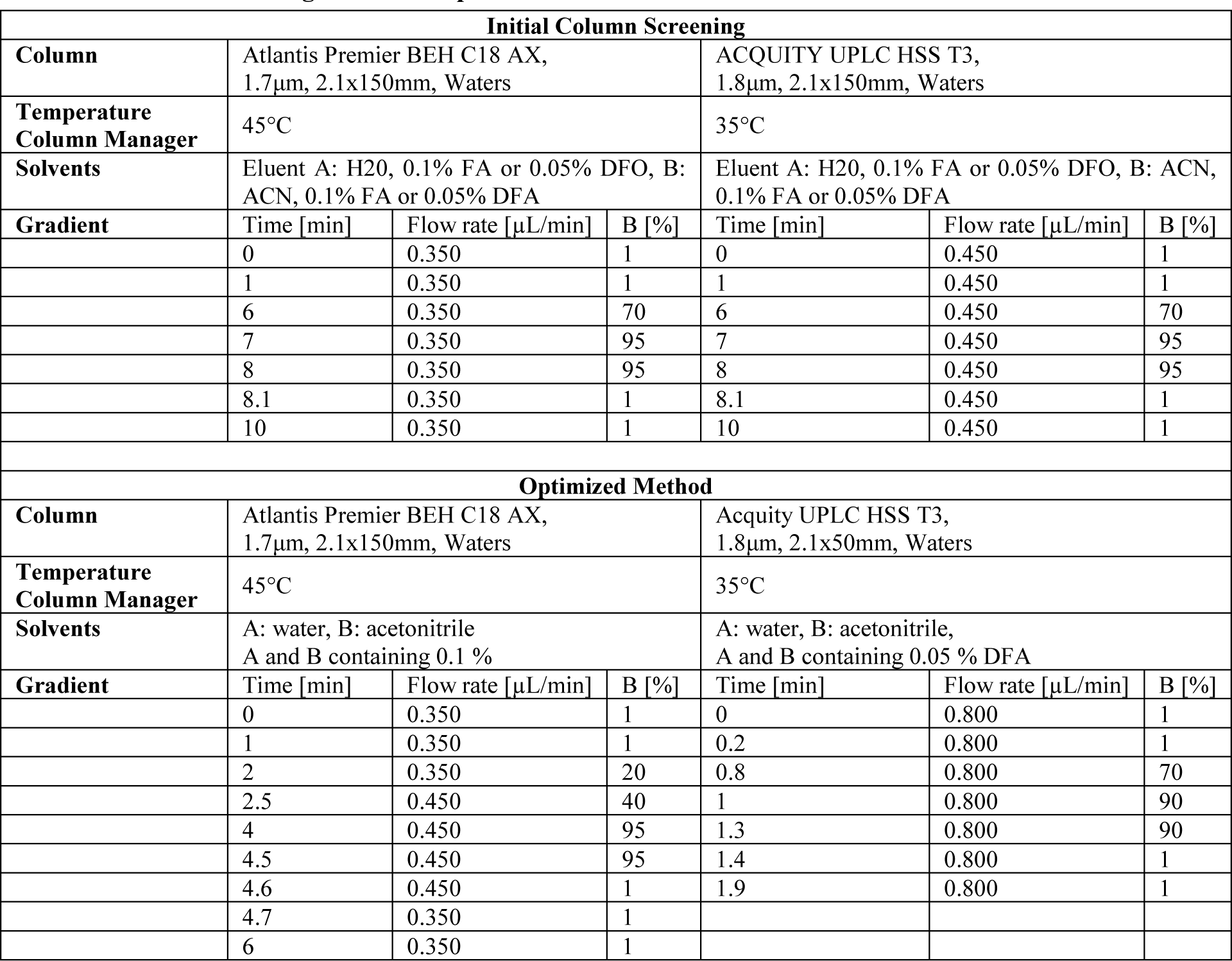
Chromatographic parameters used for the separation of amino acids and tryptophan metabolites in the initial column screening and in the optimized method.

| Initial Column Screening |  |  |  |  |  |  |
| --- | --- | --- | --- | --- | --- | --- |
| **Column** | Atlantis Premier BEH C18 AX,<br>1.7µm, 2.1x150mm, Waters |  |  | ACQUITY UPLC HSS T3,<br>1.8µm, 2.1x150mm, Waters |  |  |
| **Temperature**<br>**Column Manager** | 45°C |  |  | 35°C |  |  |
| **Solvents** | Eluent A: H2O, 0.1% FA or 0.05% DFO, B: ACN, 0.1% FA or 0.05% DFA |  |  | Eluent A: H2O, 0.1% FA or 0.05% DFO, B: ACN, 0.1% FA or 0.05% DFA |  |  |
| **Gradient** | Time [min] | Flow rate [µL/min] | B [%] | Time [min] | Flow rate [µL/min] | B [%] |
|  | 0 | 0.350 | 1 | 0 | 0.450 | 1 |
|  | 1 | 0.350 | 1 | 1 | 0.450 | 1 |
|  | 6 | 0.350 | 70 | 6 | 0.450 | 70 |
|  | 7 | 0.350 | 95 | 7 | 0.450 | 95 |
|  | 8 | 0.350 | 95 | 8 | 0.450 | 95 |
|  | 8.1 | 0.350 | 1 | 8.1 | 0.450 | 1 |
|  | 10 | 0.350 | 1 | 10 | 0.450 | 1 |
| Optimized Method |  |  |  |  |  |  |
| **Column** | Atlantis Premier BEH C18 AX,<br>1.7µm, 2.1x150mm, Waters |  |  | Acquity UPLC HSS T3,<br>1.8µm, 2.1x50mm, Waters |  |  |
| **Temperature**<br>**Column Manager** | 45°C |  |  | 35°C |  |  |
| **Solvents** | A: water, B: acetonitrile<br>A and B containing 0.1 % |  |  | A: water, B: acetonitrile,<br>A and B containing 0.05 % DFA |  |  |
| **Gradient** | Time [min] | Flow rate [µL/min] | B [%] | Time [min] | Flow rate [µL/min] | B [%] |
|  | 0 | 0.350 | 1 | 0 | 0.800 | 1 |
|  | 1 | 0.350 | 1 | 0.2 | 0.800 | 1 |
|  | 2 | 0.350 | 20 | 0.8 | 0.800 | 70 |
|  | 2.5 | 0.450 | 40 | 1 | 0.800 | 90 |
|  | 4 | 0.450 | 95 | 1.3 | 0.800 | 90 |
|  | 4.5 | 0.450 | 95 | 1.4 | 0.800 | 1 |
|  | 4.6 | 0.450 | 1 | 1.9 | 0.800 | 1 |
|  | 4.7 | 0.350 | 1 |  |  |  |
|  | 6 | 0.350 | 1 |  |  |  |

The UPLC system was coupled to a triple quadrupole (TQ) mass spectrometer (Xevo™-TQ-XS Mass, Waters, Wilmslow, UK) via a Z-spray electrospray source. The TQ-MS was operated in positive ion mode using multiple reaction monitoring (MRM) and unit-mass resolution. Cone voltages, collision energies, and MRM transitions were optimized for each metabolite using IntelliStart™ (MassLynx 4.2, Waters) and optimized parameters were manually checked (Table 2). For all measurements, the capillary voltage was set to 0.5 kV and the following gas flows were used: desolvation gas: 1000 L/hour, cone gas: 150 L/hour, nebulizer gas: 7.0 bar. The desolvation temperature was 500°C for measurements at flow rates between 350 µL/min and 450 µL/min, and 600°C at flow rates of 800 µL/min. For quantification, the raw files were processed using MS Quan in Waters_connect™ (Waters, version 1.7.0.7) using the default parameters for the processing method and with smoothing enabled (width=1, iterations=1). Absolute concentrations were determined using the sample response ratio (peak area 12C/peak area 13C), the response factor and the slope of the external calibration series. The data were further processed for statistical analysis and data visualization using R (version 4.0.3) and RStudio (version 1.4.1106) ^37^.

**Table 2.** : Parameters used for multiple reaction monitoring (MRM) mass spectrometric analysis. Cone voltage in volt [V], collision energy used for fragmentation in electron volt [eV], m/z values used for MRM transition of the analyte and stable isotope labelled internal standard (ISTD).

| Metabolite | Cone [V] | CE [eV] | MRM analyte [m/z] | MRM ISTD [m/z] |
| --- | --- | --- | --- | --- |
| 3-hydroxykynurenine (3OHKYN) | 24 | 16 | 225 > 110 | 229 > 110 |
| Alanine (ALA) | 14 | 10 | 90 > 44 | 94 > 47 |
| Arginine (ARG) | 8 | 22 | 175 > 70 | 185 > 75 |
| Asparagine (ASN) | 6 | 14 | 133 > 74 | 139 > 77 |
| Aspartate (ASP) | 4 | 8 | 134 > 88 | 139 > 92 |
| Glutamine (GLN) | 2 | 14 | 147 > 84 | 154 > 90 |
| Glutamate (GLU) | 8 | 12 | 148 > 102 | 154 > 107 |
| Glycine (GLY) | 18 | 6 | 76 > 30 | 79 > 32 |
| Histidine (HIS) | 8 | 12 | 156 > 110 | 165 > 118 |
| Indole-3-acetic acid (I3AA) | 26 | 12 | 176 > 130 | 182 > 109 |
| Isoleucine (ILE) | 20 | 16 | 132 > 69 | 139 > 74 |
| Kynurenin (KYN) | 20 | 12 | 209 > 94 | 219 > 100 |
| Kynurenic acid (KYNAC) | 20 | 30 | 190 > 116 | 196 > 122 |
| Leucine (LEU) | 20 | 12 | 132 > 86 | 139 > 92 |
| Lysine (LYS) | 8 | 16 | 147 > 84 | 155 > 90 |
| Methionine (MET) | 18 | 14 | 150 > 56 | 156 > 60 |
| Nicotinic acid (NICAC) | 20 | 18 | 123 > 80 | 130 > 85 |
| Nicotinamide (NICAM) | 20 | 20 | 124 > 80 | 129 > 85 |
| Ornithine (ORN) | 2 | 14 | 133 > 70 | 138 > 74 |
| Phenylalanine (PHE) | 20 | 10 | 166 > 120 | 176 > 129 |
| Proline (PRO) | 10 | 8 | 116 > 70 | 122 > 75 |
| Serine (SER) | 12 | 10 | 106 > 60 | 110 > 63 |
| Threonine (THR) | 2 | 20 | 120 > 56 | 120 > 60 |
| Tryptophan (TRP) | 14 | 18 | 205 > 146 | 218 > 156 |
| Tyrosine (TYR) | 14 | 28 | 182 > 91 | 192 > 98 |
| Valine (VAL) | 14 | 8 | 118 > 72 | 124 > 77 |

### Method validation

The lower limit of detection (LLOD) and lower limit of quantification (LLOQ) were determined based on linear regression analysis of the six lowest concentrations of the calibration series ^38^. For each metabolite, the standard error of the regression line (sy/x) was divided by its slope and multiplied by a factor of 3 or 9 for the LLOD and the LLOQ, respectively.

Within-run (n=5 injections per concentration) and between-run (n=20 injections per concentration) accuracy and precision were determined using five different QC samples ranging from 0.195 µM to 6.25 µM per metabolite (dissolved in 0.1% FA). For accuracy, the mean absolute percentage error (MAPE) was calculated and samples greater than 15% of the nominal values for the QC concentrations, except for the LLOQ (lower 20 % of the nominal value), were highlighted according to the ICH guideline M10 for the validation of bioanalytical methods (EMA/CHMP/ICH/172948/2019). Similarily, for the precision, samples exceeding a coefficient of variation (CV) of 15% for the QC samples were reported, except for the LLOQ which should not exceed 20%.

### Extraction of amino acids and tryptophan metabolites from serum and plasma samples Manual extraction from serum

Serum samples were thawed on ice. AAs and TRP metabolites were extracted from 40 µL of human serum by adding 160 µL of extraction solvent (MeOH containing 3.125 µM AA ISTDs). Samples were incubated on a thermo mixer ((Eppendorf, Germany) for 1 minute at 600 rpm and 4°C, followed by a centrifugation (10 minutes, 16.100 x g, 4°C, MicroStar 17R centrifuge, VWR, Darmstadt, Germany) for sample clearance. The supernatants were transferred to a new reaction vial and evaporated to dryness (Concentrator Plus™, Eppendorf, Germany). The dried samples were dissolved in 400 µL of 0.1% FA, and 1 µL was injected for LC-MS analysis.

### Semi-automated extraction from serum samples

Semi-automated extraction was performed using a liquid handling pipetting robot (Andrew+, Waters). During the development and optimization of the semi-automated extraction workflow, several protocol steps, such as sample dilution and centrifugation steps, were subsequently modified and optimized as described in Workflows I to IV (Table S2). The robot settings are shown in Table S3.

#### Workflow I

Serum samples were thawed on ice and placed in a microtube domino. 40 µL of serum was transferred to a 96-well extraction plate (Eppendorf twin.tec® 96-well LoBind® plate, conical well, 250 µL, placed in a Microplate Shaker+ Domino equipped with a 96-well cooling unit at 4°C). For deproteinization and metabolite extraction, 160 µL of extraction solvent (MeOH containing 3.125 µM AA and TRP metabolite ISTDs) was added to the serum samples (1:5 dilution) and the samples were incubated for 5 minutes at 4°C on a Microplate Shaker+ (600 rpm). The 96-well plate was then manually transferred to a centrifuge (Heraeus Megafuge 1.0R, Gemini, Germany) for sample clearance (centrifugation for 10 min at 4°C, 4,300 x g). After centrifugation, 60 µL of the supernatant was transferred by the robot to a 96-well measurement plate (Waters QuanRecovery, Waters, placed in a Microplate Shaker+ Domino) to which the robot had added 60 µL of 0.1% FA. Samples were shaken at 600 rpm for 5 minutes. The 96-well plate was manually transferred to the autosampler of the U(H)PLC system for subsequent MS analysis.

#### Workflow II

In contrast to Workflow I, a 1:9 dilution was used in Workflow II. 20 µL of serum were added to a 96- well extraction plate (Eppendorf twin.tec® 96-well LoBind® plate, conical well, 250 µL, placed in Microplate Shaker+ Domino equipped with 96 well cooling unit at 4°C) and 160 µL of extraction solvent (MeOH containing 2.8125 µM AA and TRP metabolite ISTDs). Further sample processing was performed as described in Workflow I.

#### Workflow III

Workflow III used a longer centrifugation time of 20 minutes for sample clearance. 40 µL of serum were added to a 96-well extraction plate (Eppendorf twin.tec® 96-well LoBind® plate, conical well, 250 µL, placed in Microplate Shaker+ Domino equipped with 96 well cooling unit at 4°C) and 160 µL of extraction solvent (MeOH containing 3.125 µM AA and TRP metabolite ISTDs) for deproteinization and sample clearance as described in Workflow I. For sample clearance, the sample was for 20 min at 4 °C and 4.300 x g (Heraeus Megafuge 1.0R, Gemini, Germany). After centrifugation, 60 µL of the supernatant was transferred by the robot to a 96-well measurement plate (Waters QuanRecovery, Waters, placed in a Microplate Shaker+ Domino equipped with 96 well cooling unit at 4°C) to which the robot had added 60 µL of 0.1% FA. Samples were shaken at 600 rpm for 5 minutes. The 96-well plate was manually transferred to the autosampler of the U(H)PLC system for subsequent MS analysis.

#### Workflow VI

For Workflow VI, an additional sample clearance step was used. 40 µL of serum was added to a 96-well extraction plate (Eppendorf twin.tec® 96-well LoBind® plate, conical well, 250 µL, placed in Microplate Shaker+ Domino equipped with 96 well cooling unit at 4°C), followed by the addition of 160 µL extraction solvent (MeOH containing 3.125 µM AA and TRP metabolite ISTDs), and incubated for 5 min at 4°C and 600 rpm. For the first clearance step, the samples were centrifuged for 10 min at 4 °C and 4300 x g (Heraeus Megafuge 1.0R, Gemini, Germany). Then, 60 µL of the supernatant was transferred by the robot to a 96-well extraction plate (Eppendorf twin.tec® 96-well LoBind® plate, conical well, 250 µL, placed in Microplate Shaker+ Domino equipped with 96 well cooling unit at 4°C) to which the robot had added 60 µL of 0.1% FA, and the samples were incubated for 5 min at 600 rpm. A second clearance step was then performed (centrifugation at 4.300 x g for 10 min at 4 °C (Heraeus Megafuge 1.0R, Gemini, Germany)), and 60 µL of the supernatants were transferred by the robot to a 96-well measurement plate (Waters QuanRecovery, Waters, placed in a Microplate Shaker+ Domino) to which the robot had added 60 µL of 0.1% FA. Samples were shaken at 600 rpm for 5 minutes. The 96-well plate was manually transferred to the autosampler of the U(H)PLC system for subsequent MS analysis.

#### Plasma extraction

Plasma samples were thawed on ice. For extraction, 180 µL of extraction solvent (MeOH containing 2.75 µM AA and TRP metabolite ISTD) and 20 µL of the plasma sample (1:10 dilution) were used. After the first incubation step (Microplate Shaker+ Domino, 5 min, 600 rpm, 4°C), the samples were incubated for 60 min on ice. The samples were further processed as described in Workflow IV for serum extraction.

## Results and Discussion

### Development of a rapid and sensitive U(H)PLC-MRM-MS method for quantification of underivatized amino acids and tryptophan metabolites

#### U(H)PLC-MRM-MS method development

To enable sensitive quantification of 20 amino acid (AA) and 6 tryptophan (TRP) metabolites, we first optimized the multiple reaction monitoring (MRM) transition on a triple quadrupole mass spectrometer (TQ-MS) for each individual analyte. AAs and TRP metabolite standards were introduced into the analytical flow of the U(H)PLC-MS system coupled to the TQ-MS, and the cone voltages and collision voltages were tuned to optimize analyte transmission and fragmentation (Table 1). The optimized MRM transitions were used to investigate the performance of two different column chemistries for the separation of AAs and TRP metabolites. The column chemistries, a high-strength silica reversed-phase column (ACQUITY Premier HSS T3 column, hereafter referred to as RP column) ^39,40^, and a bridged ethyl siloxane/silica hybrid (BEH) mixed-mode (MM) column with reversed-phase and weak anion-exchange properties (Atlantis Premier BEH C18 AX, hereafter referred to as MM column) ^41^, were specifically designed to retain and separate polar compounds. The 26 AA and TRP metabolite standards were pooled in solvent matrix (0.1% FA) and 6.25 pmol were injected onto the MM column (1.7 µm, 2.1 x 150 mm) or the RP column (1.8 µm, 2.1 x 150 mm). The analytes were separated on both columns using the same 10 min H20-ACN gradient and mobile phase modifiers. Due to the different particle sizes, the separation on the MM column (1.7 µm) was performed at a flow rate of 350 µL/min and the separation on the RP column (1.8 µm) was performed at a flow rate of 450 µL/min. The mobile phase modifiers used were 0.1% formic acid (FA) and 0.05% difluoroacetic acid (DFA) ^42–44^. FA is one of the most commonly used mobile phase modifiers in LC-MS applications. As a stronger ion-pairing reagent, DFA showed improved peak shapes for peptides and proteins compared to FA, and an increased electrospray ionization efficiency compared to TFA ^44,45^. However, little is known about the effects of DFA as a mobile phase modifier for LC-MS analysis of small molecules including free AAs and TRP metabolites.

For the separation on the MM column, all detected metabolites showed baseline separation in their individual MRM traces, and the isobaric amino acids isoleucine (ILE) and leucine (LEU) were baseline-separated with both 0.1% FA and 0.05% DFA as modifiers (Figure S1A). The majority of the metabolites, especially smaller amino acids without aromatic ring structure, eluted in the first 1.5 min. Kynurenic acid (KYNAC) was the only metabolite that was not detected with 0.1% FA. With 0.05% DFA as mobile phase additive, retention was similar to that observed with 0.1% FA. Comparing the mean peak widths of all metabolites, DFA (mean w1/2; DFA=2.01 s) yielded slightly sharper peaks than FA (mean w1/2; FA=2.10 s) (Table S4), but the use of DFA for ILE (w1/2; DFA=2.29 s) resulted in a slightly broader peak than with FA (w1/2; FA=2.13 s). The broadening of the ILE peak using DFA (Figure S1A) could lead to problems with reproducible baseline separation of the isobaric ILE and LEU in larger studies. In addition, glutamine (GLN) and glutamate (GLU) could not be baseline-separated with DFA. Partial coelution affected the absolute quantification of GLN and GLU in MRM mode. GLN and GLU could not be separated from each other in the first mass-selective quadrupole of the TQ-MS, and collision-induced dissociation (CID) resulted in isobaric fragments for GLU and GLN using uniformly 13C- and 15N-labeled standards ^26^. As with 0.1% FA, KYNAC could not be detected with DFA. However, compared to 0.1% FA, aspartate (ASP) nicotinic acid (NICAC), and nicotinamide (NICAM) could not be detected with DFA as a mobile phase modifier, so we concluded that FA was a better modifier for the separation of AAs and TRP metabolites using the MM C18 AX column.

For the separation on the RP HSS T3 column, earlier retention times were observed for the majority of the AAs compared to the MM column with both 0.1% FA and 0.05% DFA as modifiers (Figure S1B). The isobaric amino acids ILE and LEU could not be separated with baseline resolution. The same was observed for LYS and GLN, as well as asparagine (ASN) and ASP. The partial coelution of these AAs impairs absolute quantification by MRM using uniformly 13C- and 15N-labeled standards. The precursor ions could not be separated in the first mass-selective quadrupole of TQ-MS, and the CID fragment ions were isobaric. The remaining AAs and all tryptophan metabolites, including KYNAC, that could not be detected on the MM column, were successfully detected and separated from each other. As with the MM separation, DFA provided slightly narrower half- peak widths (mean w1/2; DFA=1.93 s) than FA (mean w1/2; FA=2.06 s) (Table S4).

The results of the column screening showed that the BEH C18 AX column with 0.1% FA provided the highest coverage and separation efficiency for free AAs and TRP metabolites including uniformly 13C- and 15N-labeled ISTDs, and that only KYNAC could not be detected with the mixed-mode column.

We then evaluated the performance of the BEH C18 AX for the separation and detection of free AAs and TRP metabolites extracted from human serum using ice-cold methanol. The majority of analytes were successfully detected and separated. However, peak splitting was observed for the basic amino acids arginine (ARG) and histidine (HIS) (Figure S2A), and neither NICAC, NICAM, nor the corresponding ISTDs could be detected in human serum samples. To overcome these limitations and to detect KYNAC, an additional 2.1 x 50 mm RP HSS T3 column (1.9 µm) was installed in the column compartment of the U(H)PLC system. Samples were injected onto the short HSS T3 column and separated within 1.9 min at a flow rate of 800 µL/min using 0.1% FA or 0.05% DFA as mobile phase modifiers. ARG, HIS, NICAC, NICAM and KYNAC were detected in the human serum extracts (Figure S2B, C). Compared to the MM column, no peak splitting was observed for ARG and HIS. Comparison of the two mobile phase additives showed that the highest separation efficiency was obtained with 0.05% DFA (Figure S2C), especially ARG, HIS and KYNAC showed improved peak shape compared to 0.1% FA (Figure S2B).

From our column screening, we concluded that a dual-column U(H)PLC-MRM-MS setup consisting of a 2.1 x 150 mm BEH C18 AX column (1.7 µm) and a 2.1 x 50 mm HSS T3 column (1.8 µm) provided optimal results for the rapid analysis of the 20 free underivatized AAs and 7 TRP metabolites with a total run time of 7.9 min. The BEH C18 AX MM column was used for the analysis of 18 AAs and 4 TRP metabolites within 6 min, and the HSS T3 RP column was used for the analysis of ARG, HIS, KYNAC, NICAC and NICAM within 1.9 min. The total run time for the analysis of the 26 AAs and TRP metabolites was 7.9 min.

#### Linearity, lower limit of detection (LLOD) and lower limit of quantification (LLOQ)

To determine the linearity of the dual-column U(H)PLC-MRM-MS method, a dilution series of unlabeled metabolite standards was prepared from 0.05µM to 6.25µM. A constant amount of stable isotope labeled metabolite standards (1.25µM) was added to each sample, and the samples were measured four times. An excellent coefficient of determination (R^2^) was observed for all metabolites over the entire concentration range (Figure 1A, Figure S3). Except for LYS (R^2^ = 0.988), indole-3-acetic acid (I3AA, R^2^ = 0.989), tyrosine (TYR, R^2^ = 0.989) and GLY (R^2^ = 0.983), all metabolites showed R^2^ values higher than 0.990, highlighting the excellent linearity of our dual-column U(H)PLC-MRM-MS setup.

**Figure 1:**
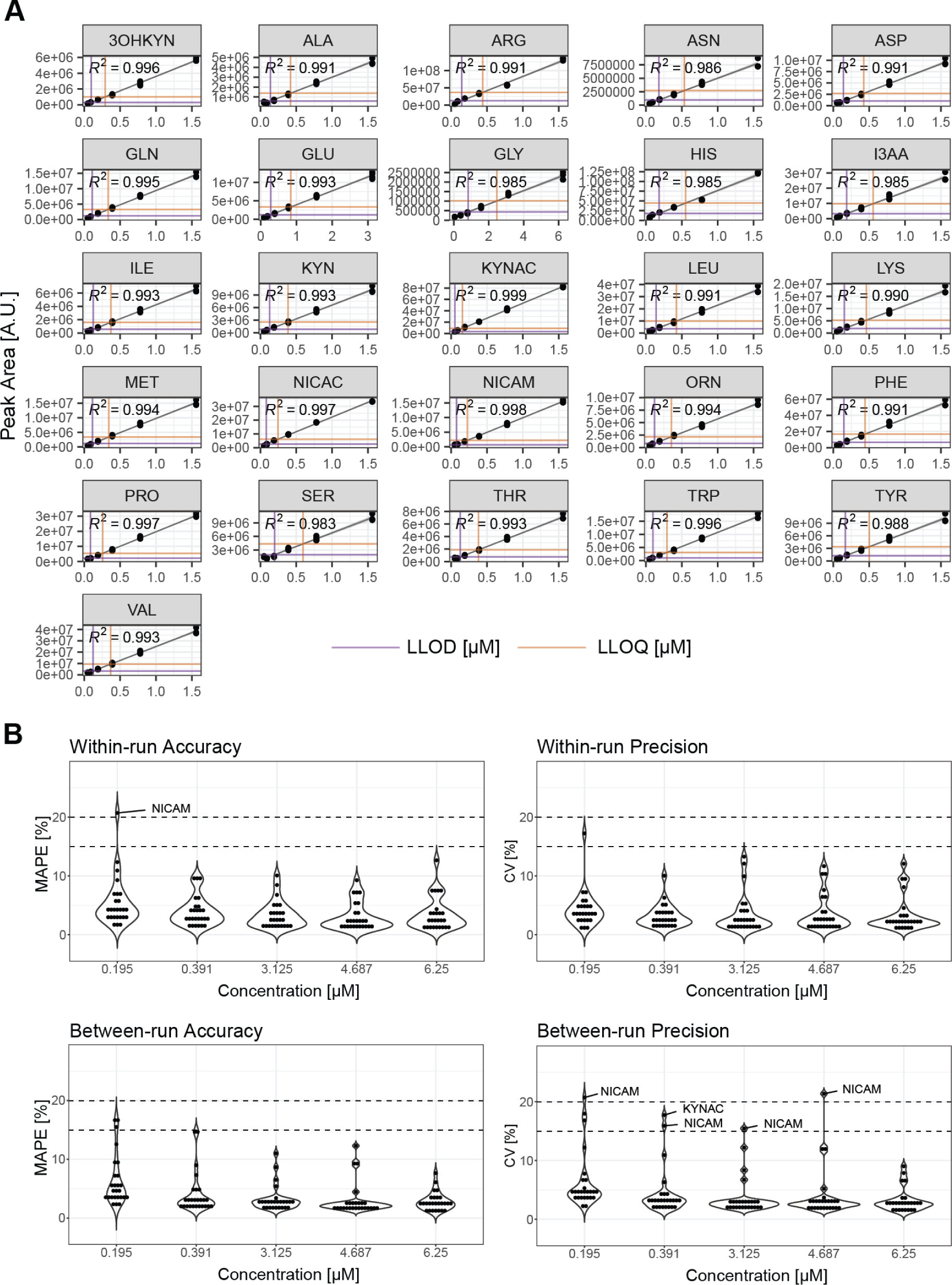
Quantification of amino acids and tryptophan metabolites using dual-column U(H)PLC-MRM-MS. A: Lower limit of detection (LLOD) and lower limit of quantification (LLOQ) for metabolite standards dissolved in the solvent matrix. Coefficient of determination (R^2^) and calculated concentrations for LLOD (purple) and LLOQ (orange) are for each metabolite (n= 4). B: Within-run (n=5) and between-run accuracy (n=20) represented by the positive amount of mean absolute percentage error (MAPE) per QC concentration, metabolites with MAPE values ≥ 20% are indicated. Within-run (n=5) and between-run (n=20) precision represented by the coefficient of variation (CV) per QC concentration, metabolites with CV values ≥ 20% at 0.195 µM or CV values ≥ 15% at all other concentrations are indicated. NICAM: Nicotinamide, KYNAC: Kynurenic acid.

To determine the sensitivity of our method for the 26 metabolites, namely the lower limit of detection (LLOD) and the lower limit of quantification (LLOQ), we used the slope of the six lowest concentrations of the dilution series (calibration curve) and the standard error of this calibration curve ^38^. For all metabolites except GLY (LLOQ: 2.46 µM), the dual-column U(H)PLC-MRM-MS method provided LLODs and LLOQs in the nanomolar range (Figure 1A, Table 3). The current gold standard for amino acid analysis in serum or plasma, namely ion exchange chromatography (IEX) followed by post-column ninhydrin derivatization and fluorescence detection, provided LLODs and LLOQs in the low µmolar range ^46^. Our dual-column U(H)PLC-MRM-MS method provided lower LLOD and LLOQ concentrations for all amino acids analyzed and was 15 times faster than the gold standard (run time: 119 min) with total a run time of 7.9 min. Choi et al. ^32^, Liu et al. ^47^ and DeArmond et al. ^33^ recently developed HPLC-MRM-MS methods based on mixed-mode chromatography using hydrophilic interaction chromatography (HILIC) and cation exchange (CEX) properties to separate and quantify free amino acids. They achieved LLOQ values in the low µmolar range with analysis times ranging from 13 min ^47^ to 15 min ^33^ and 22 min ^32^, respectively. Compared to these methods, our dual-column U(H)PLC-MRM-MS method provided lower LLOQ concentrations for the amino acids analyzed in a shorter analysis time. For TRP metabolites, we obtained similar LLOQ concentrations in the nanomolar range for the analyzed TRP metabolites compared to Desmons et al., who recently developed a 10 min HPLC-MRM-MS method for the quantification TRP metabolites but no amino acids using a biphenyl column for chromatographic separation ^18^. Taken together, these results demonstrated that our dual-column U(H)PLC-MRM-MS method provided high sensitivity suitable for the analysis of free amino acids and TRP metabolites in human serum and plasma samples.

**Table 3.** Lower limit of detection (LLOD) and lower limit of quantification (LLOQ) of the quantified metabolites.

| Metabolite | LLOD [μM] | LLOQ [μM] |
| --- | --- | --- |
| 3-hydroxykynurenine (3OHKYN) | 0.097 | 0.292 |
| Alanine (ALA) | 0.140 | 0.420 |
| Arginine (ARG) | 0.147 | 0.441 |
| Asparagine (ASN) | 0.179 | 0.536 |
| Aspartate (ASP) | 0.141 | 0.424 |
| Glutamine (GLN) | 0.111 | 0.334 |
| Glutamate (GLU) | 0.280 | 0.841 |
| Glycine (GLY) | 0.822 | 2.467 |
| Histidine (HIS) | 0.184 | 0.552 |
| Indole-3-acetic acid (I3AA) | 0.185 | 0.555 |
| Isoleucine (ILE) | 0.124 | 0.372 |
| Kynurenin (KYN) | 0.128 | 0.385 |
| Kynurenic acid (KYNAC) | 0.053 | 0.158 |
| Leucine (LEU) | 0.142 | 0.425 |
| Lysine (LYS) | 0.155 | 0.465 |
| Methionine (MET) | 0.114 | 0.342 |
| Nicotinic acid (NICAC) | 0.082 | 0.245 |
| Nicotinamide (NICAM) | 0.075 | 0.226 |
| Ornithine (ORN) | 0.121 | 0.363 |
| Phenylalanine (PHE) | 0.178 | 0.534 |
| Proline (PRO) | 0.086 | 0.257 |
| Serine (SER) | 0.198 | 0.594 |
| Threonine (THR) | 0.127 | 0.381 |
| Tryptophan (TRP) | 0.097 | 0.292 |
| Tyrosine (TYR) | 0.167 | 0.501 |
| Valine (VAL) | 0.123 | 0.368 |

#### Within-run and between-run accuracy and precision

To monitor the within-run and between-run accuracy and precision, 5 different QC samples ranging from 0.195 µM to 6.25 µM were injected multiple times over different time periods according to the ICH Guideline M10 for Validation of Bioanalytical Methods (EMA/CHMP/ICH/172948/2019). Within-run accuracy and precision were determined by measuring five replicates per concentration within an analytical run. Accuracy was within 15 % mean absolute percentage error (MAPE) for all analytes at all concentrations except NICAM (20.7%) at 0.19 µM (Figure 1B, Table S5). In terms of precision, the coefficient of variation (CV) was less than 20% for all metabolites at the lowest QC level and below 15% at all other levels. Low mean values for within-run accuracy (MAPE=4.0%) and precision (CV=3.6%) were observed over all concentrations.

To evaluate between-run accuracy and precision, 10 analytical runs were performed over 3 days. The between-run accuracy was less than 20% at 0.19 µM and less than 15% at the other four QC levels for all analytes (Figure 1B, Table S6). Similar results were obtained for the between-run precision, except for NICAM at 0.19 µM (MAPE=20.7%), 0.39 µM (MAPE=15.9%), 3.12 µM (MAPE=15.5%) and 4.7 µM (MAPE=21.4%) and KYNAC at 0.39 µM (MAPE=17.7%), all metabolites showed CV values below 20% at 0.19 µM and below 15% at the other OC levels. Furthermore, all metabolites showed good values for accuracy and precision, as reflected by low mean values for between-run accuracy (MAPE=3.9%) and precision (CV=4.4%), highlighting the excellent within-run and between-run accuracy and precision of our dual-column U(H)PLC-MRM-MS method.

In conclusion, our dual-column U(H)PLC-MRM-MS method allows the quantification of 20 amino acids and 6 TRP metabolites without derivatization within a total run time of 7.9 min. The U(H)PLC-MRM-MS methods showed excellent linearity over a high concentration range, as well as excellent within-run and between-run precision and accuracy. The high sensitivity with LLOQs in the nanomolar range except for glycine (LLOQ: 2.46 µM) and linearity should be suitable for the quantification of free AA and TRP metabolites from human serum and plasma.

#### Semi-automated extraction and sample preparation of human serum and plasma

Quantitative analysis of free amino acids and TRP metabolites in large patient cohorts requires rapid and robust analyte extraction and sample processing to ensure high reproducibility and sample quality. To achieve this, we developed a semi-automated workflow with a focus on minimizing the number of processing steps and automating as many as possible. The Andrew+ robotic liquid handling platform was used for automation. The liquid handler did not include a centrifuge module. The transfer of the sample plates to and from the centrifuge and the operation of the centrifuge had to be performed manually. For this reason, we refer to the workflow as a semi-automated workflow.

#### Manual extraction and sample preparation versus semi-automated workflow

First, a manual extraction of amino acids from human serum was compared with a semi-automated extraction workflow. Manual extraction and sample processing were performed by two experienced operators. Each operator performed three independent experiments (total: six independent experiments), and eight independent experiments were performed with the pipetting robot. For the extraction, MeOH together with stable isotope-labeled AA standards was added to the serum, followed by a sample clearance step. The supernatants were transferred to a new reaction vial containing 0.1% FA (final sample solution: 50% MeOH, 0.05% FA), and 1 µL was injected for LC-MS analysis using the MM column. As expected, the semi-automated extraction resulted in significantly less variation for both endogenous AAs and ISTDs compared to the manual extraction (Figure 2A). While all analyzed endogenous AAs and ISTDs showed CV values below 25% in the case of the semi-automated extraction, all analytes (endogenous AAs and ISTDs) showed CV values above 25% in case of the manual extraction. With the exception of TRP, the semi-automated workflow resulted in CV values less than 15% for both endogenous AAs and ISTDs. After normalization of endogenous AAs to the ISTD, all AAs except and GLN showed CV values below 15% for both the semi-automated extraction and the manual extraction. However, AA extraction using the semi-automated workflow still resulted in a significantly lower variation (Figure 2A). This result highlights the advantage of a semi-automated AA extraction workflow in terms of reproducibility compared to manual extraction and sample processing. Especially when multiple operators are responsible for sample preparation, which is a realistic scenario when large cohorts are processed manually in a laboratory.

**Figure 2:**
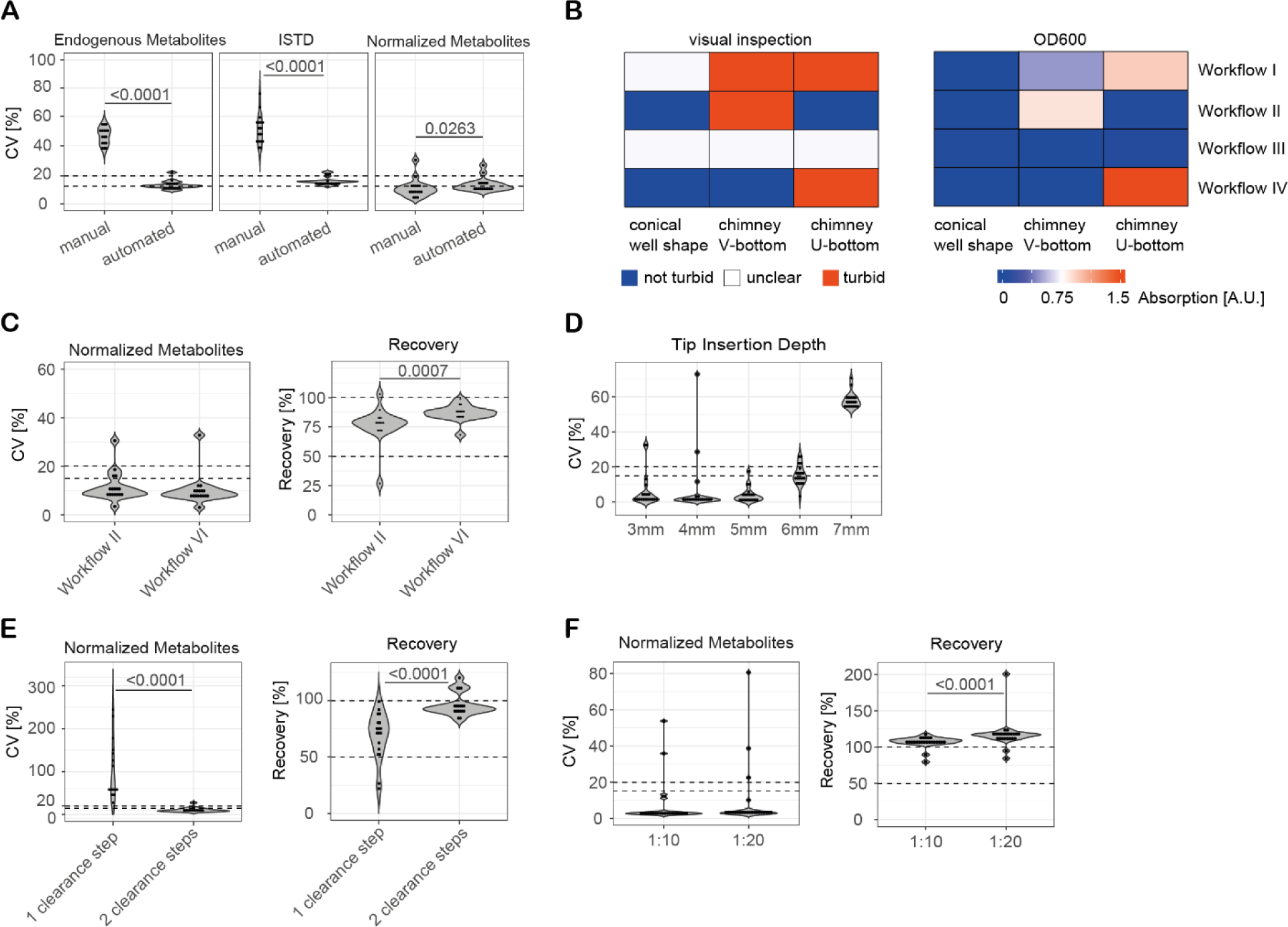
Optimization of the semi-automated extraction of amino acids and tryptophan metabolites from human serum and human plasma samples. A: Comparison of the performance of manual amino acid extraction (two operators, n= 6 independent experiments) and semi-automated amino acid extraction (n=8 independent experiments) from human serum. CV: coefficient of variation [%], ISTD: stable isotope-labeled internal standards. B: Optimization of deproteinization and sample clearance of the semi-automated extraction (n=8 independent experiments per condition). Workflow I: Sample dilution 1:5, centrifugation: 4,300 rpm, 10 min, 4°C. Workflow II: Sample dilution 1:9, centrifugation: 4,300 rpm, 10 min, 4°C. Workflow III: Sample dilution 1:5, centrifugation: 4,300 rpm, 20 min, 4°C. Workflow IV: Sample dilution 1:5, centrifugation two times: 4,300 rpm, 10 min, 4°C. Visual inspection (turbidity (red), no turbidity (blue), unclear result (white)) compared to spectrophotometric analysis of optical density at 600 nm (OD 600) using a heat map (absorption) to distinguish between samples with a low turbidity (blue color) and high turbidity (red color). C: Dual-column U(H)PLC-MRM-MS analysis of samples extracted from human serum using Workflow II or Workflow IV (see description above), CV: coefficient of variation [%] and recovery % (n=8 independent experiments per workflow). D: Evaluation of the tip insertion depth of the robotic liquid handling platform, CV: coefficient of variation [%] of the added stable isotope-labeled internal standards (ISTD) (n=3). E: Semi-automated metabolite extraction from human plasma samples using one or two sample clearance steps (centrifugation: 4,300 rpm, 10 min). CV: coefficient of variation [%], recovery [%] (n=8 independent experiments). F: Semi-automated metabolite extraction from human plasma samples using a 1:10 sample dilution or 1:20 sample dilution, CV: coefficient of variation [%], recovery [%] (n=8 independent experiments). For statistical analysis, p-values were calculated using the Mann-Whitney U test. Dotted lines represent CV value of 15% and 20% or a recovery of 50% and 100%, respectively.

#### Optimization of the semi-automated extraction and sample preparation workflow for serum samples

During the initial semi-automated extractions, an insufficient deproteinization and sample clearance was observed. As a result, precipitates were observed in the reaction vials of the samples subjected to LC-MS analysis, and the pipettes were partially clogged, resulting in variations in the volumes to be transferred during sample processing. To improve deproteinization and sample clearance, we evaluated three different well plate shapes, namely U-bottom well plates, V-bottom well plates, and conical well plates, for their ability to produce stable protein pellets. The semi-automated extraction workflow was also modified to increase deproteinization and sample clearance. In addition to the initial extraction workflow using a 1:5 sample dilution and one clearance step (centrifugation: 4,300 rpm, 10 min) (Workflow I), a 1:9 sample dilution was used (Workflow II), the centrifugation time of the clearance step was increased to 20 min (Workflow III), and a second clearance step was added (Workflow IV, total of two clearance steps at 4,300 rpm for 10 min). The deproteinization and clearance performance was evaluated by visual inspection and spectroscopic analysis of the cleared samples (Figure 2B). For visual inspection, the following criteria were used: sample was turbid (Figure 2B, red), sample was not turbid (Figure 2B, blue), it was not possible to determine if the sample was turbid and not (unclear result, Figure 2B, white). For the spectroscopic analysis, the optical density of the cleared samples was measured at a wavelength of 600 nm, OD600 is a measure of the light scattering in a solution caused, for example, by protein aggregates ^48^. The results of the visual analysis were compared and correlated with the results of the spectroscopic analysis and presented in a heatmap to distinguish between samples with a low turbidity (blue) and high turbidity (red) (Figure 2B). Workflow II (reduced sample volume and high sample dilution (1:9)) and Workflow IV (addition of a second clearance step) in combination with the conical shaped 96- well plate provided the most efficient deproteinization and sample clearance, as indicated by the lowest turbidity, and were therefore selected for further evaluation using the dual-column U(H)PLC-MRM-MS method.

Workflow IV, using a 1:5 sample dilution for extraction combined with two clearance steps, resulted in a higher reproducibility (mean CV over all analytes = 10.1+/-6.2%) and significantly higher recovery (mean recovery over all analytes = 86.9+/-6.2%) than Workflow II, using a 1:9 sample dilution for extraction combined with one clearance step (mean CV over all analytes = 11.6+/-6.1%, mean recovery over all analytes = 76.7+/-14.9%) (Figure 2C, Table S7). With the exception of threonine (CV= 32.8%, recovery: 68.2), all analytes showed CV values less than 15% and a recovery greater than 80%.

Finally, the insertion depth was evaluated and optimized. If the tip is inserted too deep into the well, the pellet may be disturbed and precipitates may be transferred along with the supernatant, whereas if the tip is not inserted deep enough, the transfer may be incomplete. Our results showed that an insertion depth of 5 mm measured from the bottom of the well resulted in CV < 20% values for all analytes and provided the most robust results (mean CV over all analytes = 3.8+/-4.5%) (Figure 2D).

The optimized semi-automated workflow for metabolite extraction from serum samples consisted of a 1:5 sample dilution combined with two clearance steps and a tip insertion depth of 5 mm (Scheme 1).

**Scheme 1:**
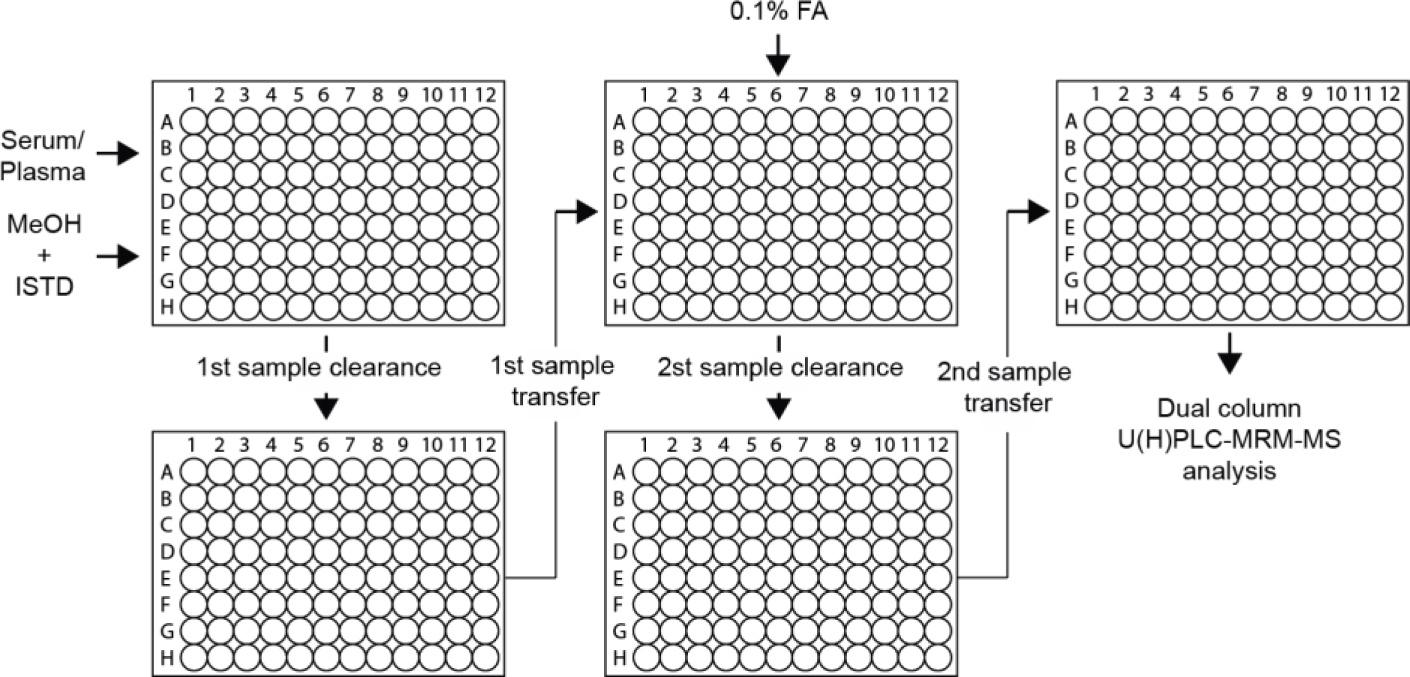
Semi-automated workflow for the extraction of amino acids and tryptophan metabolites using a robotic liquid handling platform. Sample dilution: 1:5 for serum samples, 1:10 for plasma samples. Extraction solvent: 100% methanol (MeOH) containing stable isotope-labeled internal standards (ISTD). Two sample clearance and sample transfer steps are performed, and the samples can be directly analyzed by dual-column ultrahigh performance liquid chromatography-multiple reaction monitoring mass spectrometry (U(H)PLC-MRM-MS).

#### Optimization of the semi-automated extraction and sample preparation workflow for plasma samples

For clinical and preclinical studies, plasma samples are often available instead of serum samples. While serum is the fluid that remains after the blood has clotted, plasma is the fluid that remains when clotting is prevented by the addition of an anticoagulant. Thus, the composition of serum and plasma is different. We therefore investigated whether our semi-automated workflow could also be applied to plasma samples. Since we had already identified critical steps in the workflow that needed to be optimized, we optimized the sample dilution factor and the number of clearance steps. As with the serum samples, the highest reproducibility (mean CV over all analytes = 10.4+/-5.1%) and recovery (mean recovery over all analytes = 96.4%) were achieved with two clearance steps for the plasma samples (Figure 2E, Table S8). For the dilution factor, the highest reproducibility (mean CV over all analytes = 4.7+/-7.2%) and good recovery (mean recovery over all analytes = 106%) were obtained with using a 1:10 sample dilution. With the exception of KYNAC (CV=35.8%), all analytes showed CV values below 15% (Figure 2F, Table S9).

#### Recovery

To evaluate the recovery of AAs and TRP metabolites from serum sample and plasma samples using the optimized semi-automated extraction workflows (Scheme 1), stable isotope-labeled ISTDs were added to either seven serum samples or plasma samples at the beginning or the end of the extraction. Recovery in percent was calculated by dividing the response of the stable isotope-labeled ISTD added at the beginning of the extraction by the stable isotope-labeled ISTDs added at the end of the extraction and multiplied by 100. All AAs and TRP metabolites showed high recoveries between 80 and 110% in plasma (mean recovery: 94.7%) and in serum (mean recovery: 100.9%) (Figure 3A).

**Figure 3:**
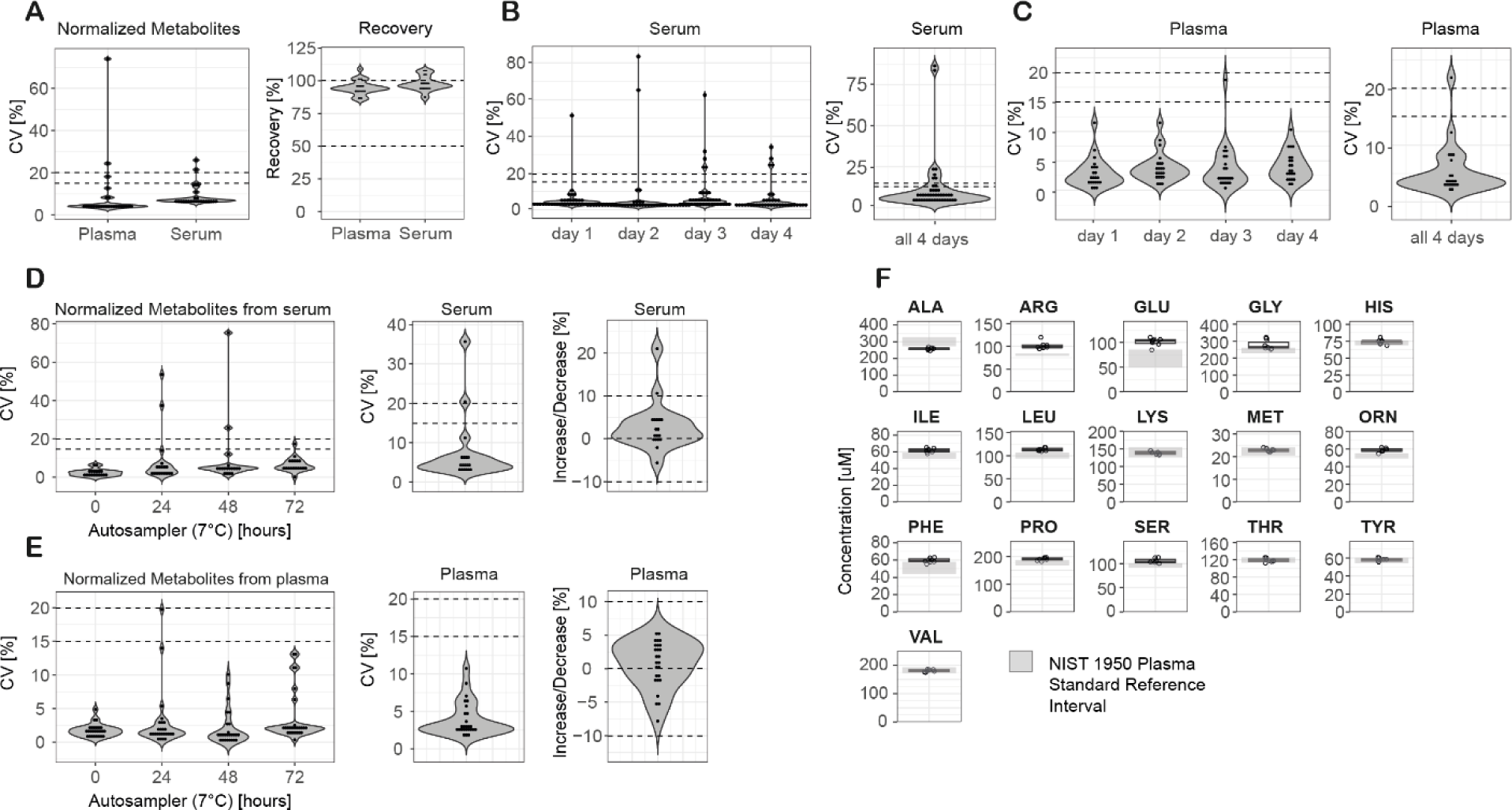
Validation of the semi-automated extraction of amino acids and tryptophan metabolites from human serum and human plasma samples. A: Schematic of the optimized semi-automated extraction workflow. B: Reproducibility (coefficient of variation (CV) [%)]) and recovery [%] of the semi-automated extraction of amino acids and tryptophan metabolites from human serum and human plasma (n= 7 independent experiments). C: Inter- day stability of the semi-automated extraction from human serum samples. CV: coefficient of variation [%] for each day of extraction (n= 8 independent experiments per day) and over all 4 days. D: Inter-day stability of the semi- automated extraction from human plasma samples. CV: coefficient of variation [%] for each day of extraction (n=3 independent experiments per day) and over all 4 days. E: Autosampler stability (7°C) of the analytes extracted from human serum, CV: coefficient of variation [%] for individual time points (n= 3 independent experiments per time point), CV% over all time points and percentage increase/decrease within 72 hours. F: Autosampler stability (7°C) of the analytes extracted from human plasma, CV: coefficient of variation [%] for individual time points (n= 3 independent experiments per time point), CV% over all time points and percentage increase/decrease within 72 hours. G: Accuracy of the absolute quantifications of amino acids extracted from the NIST SRM 1950 plasma sample using the optimized semi-automated workflow (n= 7 independent experiments). The grey rectangle represents the confidence interval and the solid horizontal line represents NIST SRM 1950 reference concentration in µmol/L. Dotted lines represent CV values of 15% and 20%, recovery of 50% and 100%, and +/- 10% analyte increase/decrease, respectively.

#### Intra-assay variability and inter-assay precision

The intra-assay variability and inter-assay precision of the semi-automated extraction were evaluated as proposed by the European Bioanalysis Forum for the analysis of metabolites in serum samples and plasma samples with quality assurance criteria of +/-20% for coefficient of variation and mean bias ^49^. Seven plasma samples and 7 serum samples were extracted for intra-assay variability. The semi-automated workflow provided high reproducibility for both serum samples and plasma samples (Figure 3A, Table S10). Except for ASP (CV= 24%) and NICAC (CV=74%) in plasma samples, and 3OHKYN (CV= 26%) and ARG (CV= 21%) in serum samples, all analytes showed CV values less than 20% and a mean intra-assay variability of 9.2% for serum samples and 8.8% for plasma samples. Inter-assay precision was calculated from the extraction of serum samples and plasma samples on four independent days. For the serum samples, the mean coefficient of variation ranged from 6.5 to 8.3% for the different days of extraction and 10.7% for all 4 days (Figure 3B, Table S11). Except for NICAC (day 1: CV= 51%, day 2: CV= 83%, day 3: CV= 62%), SER (day 3: CV= 22%, day 4: CV= 23%), 3OHKYN (day 3: CV= 24%) and GLY (day 4: CV= 34%), all analytes showed CV values less than 20%. For plasma samples, all analytes showed CV values less than 20% (Figure 3C, Table S11). The mean coefficient of variation for each day was between 3.2 to 4.8% and 5.5% over all 4 days.

#### Autosampler stability

Autosampler stability was investigated to determine how long the analytes were stable during LC-MS analysis. AAs and TRP metabolites were extracted from serum or plasma (n= 3 independent experiments), and the extracts, dissolved in LC-MS running buffer, were stored at -80°C. Samples were thawed once and placed in the autosampler to stay for 72 hours, 48 hours, 24 hours, and 0 hours, respectively, and the samples were injected together on the same day for dual-column U(H)PLC-MRM-MS analysis. Good within-day reproducibility was observed for both serum samples (mean CV values ranging from 2.3% to 9.7%) (Figure 3D, Table S12) and plasma samples (mean CV values ranging from 1.8% to 3.8%) (Figure 3E, Table S13). For all samples over the four days, a mean CV value of 7.4% was observed for serum samples and a mean CV value of 4.0% for plasma samples. To determine analyte stability over 72 hours, a percentage decrease or increase was calculated by subtracting the peak areas of the stable isotope-labeled ISTDs resulting from the time 0 hours measurement from the peak areas measured at time 72 hours, divided by the peak areas measured at time 0 hours, and multiplied by 100, as previously described by Gray et al ^50^. For plasma samples, the percentage decrease or increase was less than 10% for all analytes analyzed with a mean value of 0.5%. For serum samples, all analytes except LEU (10.5%) showed a percentage decrease or increase of less than 10% with a mean value of 2.9%. Therefore, the analytes can be considered stable in the autosampler for at least 72 hours.

### Analysis of NIST SRM 1950 reference plasma sample

To evaluate the accuracy of the semi-automated extraction workflow and the dual-column U(H)PLC- MRM-MS method, AAs and TRP metabolites were extracted from commercially available reference plasma samples (NIST SRM 1950). Seven independent experiments were performed and the AA concentrations determined were compared with the certified reference values of the NIST SRM 1950 sample (Figure 3F, Table S14). For TRP and the other TRP metabolites, no certified reference values exist for the NIST SRM 1950 plasma sample. Comparison showed that our method provided mean absolute percentage errors (MAPE) of less than 15% for 13 of the 16 analytes (Table S14). The mean MAPE over all AAs was 6.9%, and more than 50% of the analytes showed a bias of less than 10% from the reference values. The accuracy of our method was comparable to results previously published by Gray et al ^51^ and Thompson et al ^52^ evaluating 12 AAs and 13 AAs, respectively, from NIST SRM 1950 plasma samples.

### Automated extraction and analysis of serum samples from a prostate cancer study

Prostate cancer is the second most common cancer in men ^53,54^. Prostate-specific antigen (PSA) is used to screen for prostate cancer in blood samples from symptom-free men. However, PSA is also produced in non-malignant prostate gland cells, and blood PSA levels can also be elevated in non-prostate cancer conditions, including prostate infection, enlarged prostate, and due to therapies such as testosterone treatment ^55^. There is an urgent need to identify and validate new biomarker candidates and/or biomarker candidate panels because PSA screening is prone to high false-positive rates ^56^. Recent in vitro and in vivo studies in mice and humans have shown that prostate cancer may be associated with alterations in AA metabolism ^57,58^. Therefore, we used our developed semi-automated extraction workflow and dual-column U(H)PLC-MRM-MS method to analyze free AAs and TRP metabolites in serum samples from a prostate cancer study. The prostate cancer study consisted of 5 patient groups (active surveillance (AS), brachytherapy, hormone therapy, combined radiotherapy and hormone therapy, or prostatectomy) and samples from 2 control groups, healthy controls and patients with an enlarged prostate, but no cancer (referred to as the no-cancer group). AAs and TRP metabolites were extracted from 20 µL of serum. Although significant differences were observed between all seven groups for TYR (p=0.022, analysis of variation (ANOVA)), ORN (p=0.026, ANOVA) and MET (p=0.049, ANOVA) (Table S15), we chose to focus on the distinction between healthy controls and patients on active surveillance for diagnostic relevance, as no cancer treatment is provided during active surveillance. Patients on active surveillance showed lower levels of the branched-chain amino acids LEU and ILE, MET, and higher levels of ORN compared to the healthy controls (Figure 4A). Differences between most metabolites were less pronounced, but high deviations in ARG, GLU and THR levels were observed in patients on active surveillance compared to healthy controls. With the exception of TRP, which tended to be lower in patients on active surveillance, the levels of the other TRP metabolites were below the LLOQ values. Partial least squares discriminant analysis (PLS-DA) was able to successfully discriminate between the two groups based on AA levels along the first two X variates, despite the small cohort size and variation in some of the AA levels in the active surveillance cohort (Figure 4B). Dereziński et al. ^59^ and Miyagi et al. ^60^, who analyzed AA by LC-MS using pre-column derivatization from serum and plasma, respectively, observed lower levels of most AAs in prostate cancer patients, including ILE, LEU, and MET. Consistent with their findings, we also observed a trend toward lower levels of most AAs in the prostate cancer patient cohorts. In our study, the cohorts were relatively small, so that the effects observed by Dereziński et al. ^59^ and Miyagi et al. ^60^ can only be reproduced as a trend. In addition, it is not clear from the cited comparative studies which treatment the patients received. Overall, we can conclude that despite the small cohort size and sample volume, our setup was able to monitor differences in amino acid concentrations in a prostate cancer study, and it must be emphasized that both our study and the studies by Dereziński et al. ^59^ and Miyagi et al. ^60^ show that amino acid analysis from serum and plasma has potential for prostate diagnostics.

**Figure 4.**
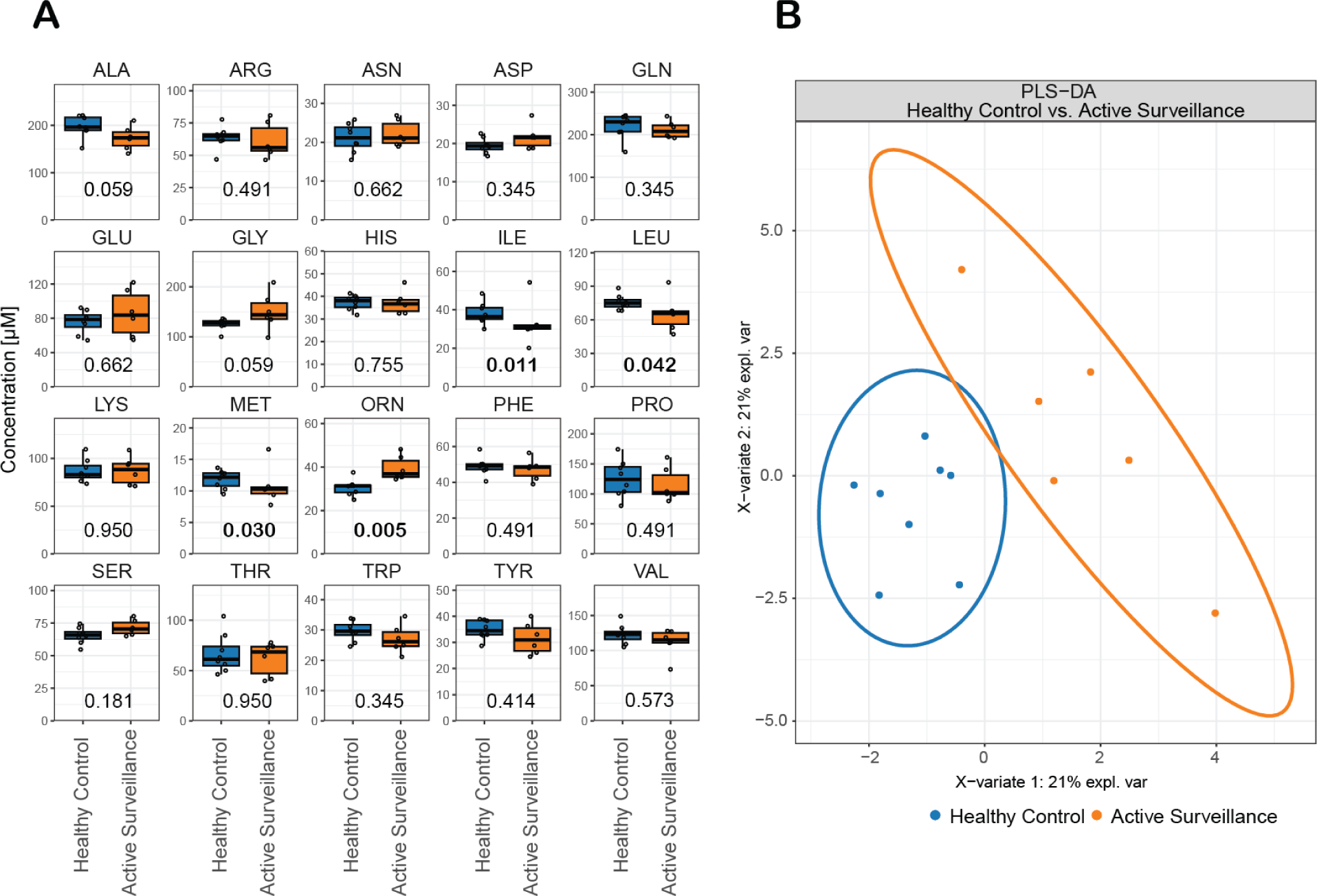
: Analysis of amino acids extracted from human serum of healthy controls (n=8) and patients on active surveillance (n=6). A: Absolute concentrations of amino acids extracted from human serum samples extracted by the semi-automated extraction workflow and analyzed by dual-column U(H)PLC-MRM-MS. Statistical analysis: Mann-Whitney U test, amino acids with p-values lower than 0.05 are highlighted in bold. B: Partial least square discriminant analysis (PLS-DA) of healthy controls and patients on active surveillance based on the quantified amino acid levels.

## Conclusion

A semi-automated workflow for the extraction of free amino acids and TRP metabolites from serum and plasma using a robotic liquid handling platform, and a dual-column U(H)PLC-MRM-MS method for the rapid absolute quantification of 20 amino acids and 6 TRP metabolites were developed and validated. The semi-automated workflow using methanol for deproteinization, two clearance steps, and conical well plates required only 20 µL of human plasma or serum respectively. The method was designed for use in a 96-well plate format with a direct interface to the autosampler of the U(H)PLC-MS system. Samples extracted in 96-well plates can be transferred directly to the U(H)PLC autosampler and analyzed by mass spectrometry without further SPE, evaporation and/or buffer exchange, and the dual-column U(H)PLC- MRM-MS method provides absolute metabolite quantification in only 7.9 minutes. The use of two different columns and ion-pairing reagents, an RP-AEX mixed-mode column with formic acid and a high-strength silica reversed-phase column with difluoroacetic acid as mobile phase additive, significantly improved the separation of isobaric free amino acids, tryptophan metabolites and, in particular, the peak shapes of basic amino acids in both serum and plasma samples. The dual-column U(H)PLC-MRM assay was found to be robust, reproducible, specific and sensitive enough for the quantification of free amino acids and TRP metabolites in plasma and serum. With LLOQs in the nanomolar range for all analytes except for GLY (LLOQ: 2.46 µM), our assay was more sensitive and 15 times faster than the gold standard for amino acid quantification, IEX, followed by post-column ninhydrin derivatization and fluorescence detection ^46^. Compared to other LC-MS assays for the quantification of underivatized amino acids ^32,33,47^ and TRP metabolites ^18^, our dual-column U(H)PLC-MRM-MS method yielded lower LLOQs for the amino acids in a 2-3 times shorter analysis time, and similar LLOQs for TRP metabolites. The semi-automated extraction in a 96-well plate format and the short LC-MS analysis time (7.9 min) make our assay ideal for the analysis of large numbers of samples in clinical and epidemiological population studies. The method has been applied to a human prostate cancer study and has been shown to discriminate between treatment regimens and to identify amino acids responsible for the statistical separation of the patient groups.

## Acknowledgment

M.K. thanks the University of Innsbruck (Project No. 316826) and the Tyrolian Research Fund (Project No. 18903) for financial support. K.T. acknowledges support from the MESI-STRAT project (European Union Horizon 2020 Research and Innovation Program, Grant Agreement No. 754688) and PARC project (funded by the European Union’s Horizon Europe Research and Innovation Program under Grant Agreement No. 101057014).

## Author Information

Corresponding author Marcel Kwiatkowski

## Author Contributions

\# Tobias Kipura and Madlen Hotze contributed equally.

## Notes

The authors declare no conflicts of interest other than those arising from employment.

