## Supplemental Figures and Tables for "Automated liquid handling extraction and rapid quantification of underivatized amino acids and tryptophan metabolites from human serum and plasma using dual-column U(H)PLC-MRM-MS and its application to prostate cancer study"

### Table of content

**Figure S1:** Chromatographic separation of amino acids (AAs) and tryptophan (TRP) metabolites **P.2**
**Figure S2:** Chromatographic separation of the basic amino acid arginine and histidine, and the tryptophan metabolites kynurenic acid, nicotinic acid and nicotinamide extracted from human serum sample. **P. 3**
**Figure S3:** Coefficient of determination (R2) of the metabolites analyzed using dual-column U(H)PLC-MRM-MS. **P. 4**
**Table S1:** Stable-isotope labeled canonical and non-canonical amino acids and tryptophan metabolites. **P.5**
**Table S2:** Parameters used for the different extraction workflows to optimize the semi-automated extraction workflow. **P. 6**
**Table S3:** Overview of optimized pipetting settings used for semi-automated extraction of amino acids and tryptophan metabolites using the robotic liquid handling platform (Andrew+). **P. 7**
**Table S4:** Peak widths at half height (w1/2) of amino acids and tryptophan metabolites. **P. 8**
**Table S5:** Within-run accuracy and within-run precision. **P. 9**
**Table S6:** Between-run accuracy and between-run precision. **P. 10**
**Table S7:** Optimization of the semi-automated extraction and sample preparation workflow for the analysis of metabolites in human serum samples. **P. 11**
**Table S8:** Optimization of the semi-automated extraction and sample preparation workflow for the analysis of metabolites in human plasma samples. **P. 12**
**Table S9:** Optimization of the semi-automated extraction and sample processing workflow for the analysis of metabolites in human plasma samples. **P. 13**
**Table S10:** Intra-assay variability of the optimized semi-automated workflow. **P. 14**
**Table S11:** Inter-assay precision of the optimized semi-automated workflow. **P. 15**
**Table S12:** Autosampler stability of the metabolites extracted from serum over 72 hours. **P. 17**
**Table S13:** Autosampler stability of the metabolites extracted from plasma over 72 hours. **P. 18**
**Table S14:** Quantification of amino acids and tryptophan metabolites extracted from reference plasma samples. **P. 19**
**Table S15:** Analysis of variances (ANOVA) for prostate cancer study. **P. 21**

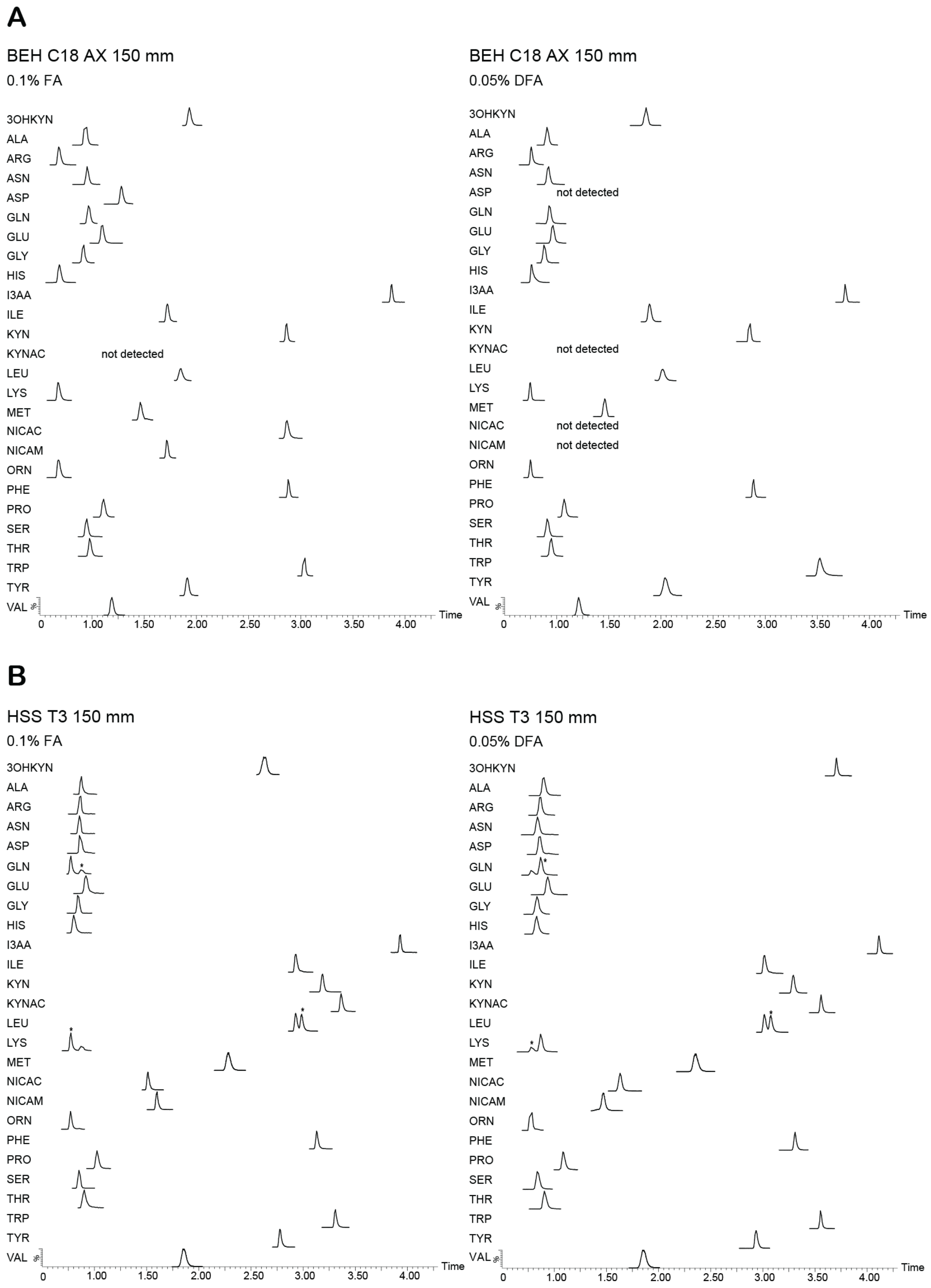


**Figure S1: Chromatographic separation of amino acids (AAs) and tryptophan (TRP) metabolites.** AAs and TRP metabolites were separated on a mixed-mode BEH C18 AX column (A) and a high-strength silica HSS-T3 reversed-phase column (B) using formic acid (FA) or difluoroacetic acid (DFA) as mobile phase additive. *: The MRM channels of LYS and GLN (both $[M+H]^+$: 147), and ILE and LEU (both $[M+H]^+$: 132) show two peaks due to the same intact mass selected in the first mass selective quadrupole and the same mass of the fragment selected in the second mass selective quadrupole. The peak of the indicated analyte is marked with an asterisk (*).

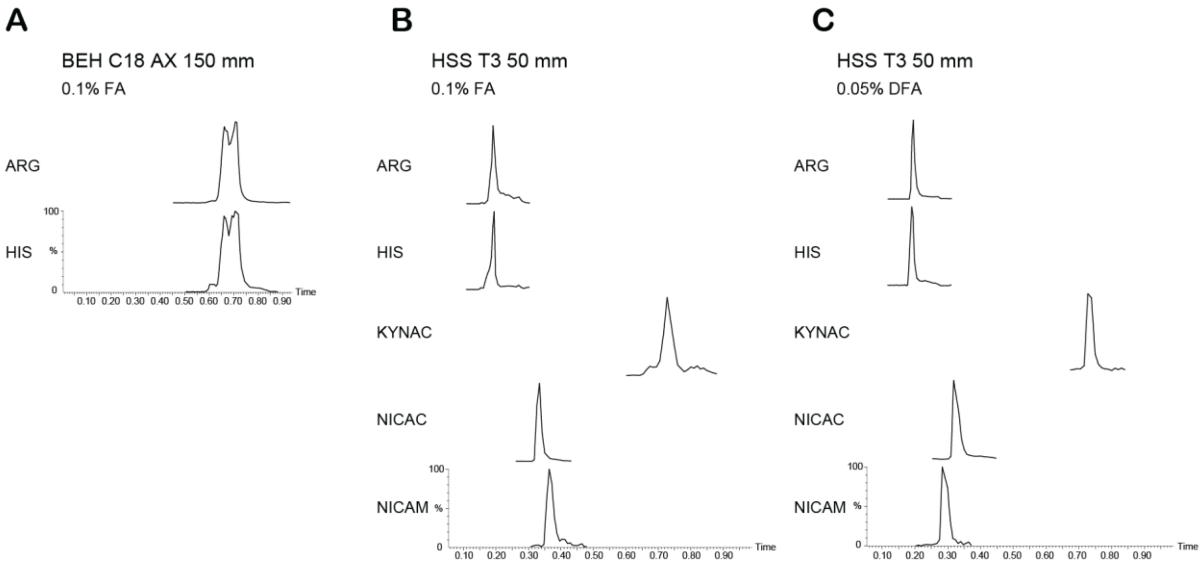


**Figure S2: Chromatographic separation of the basic arginine and histidine, and the tryptophan metabolites kynurenic acid, nicotinic acid and nicotinamide extracted from human serum sample.** A: Chromatograms of ARG and HIS separated on the BEH C18 AX mixed-mode column (2.1x150 mm, 1.7 µm) using 0.1% formic acid (FA) as mobile phase additive. KYNAC, NICAC and NICAM were not detected. B, C: Chromatograms of ARG, HIS, KYNAC, NICAC, NICAM separated on the HSS T3 reversed-phase column (2.1x50 mm, 1.8 µm) with 0.1% formic acid (FA) (B) or 0.05% difluoroacetic acid (DFA) (C) as mobile phase additive. ARG: Arginine, HIS: Histidine, KYNAC: Kynurenic acid, NICAC: Nicotinic acid, NICAM: Nicotinamide.

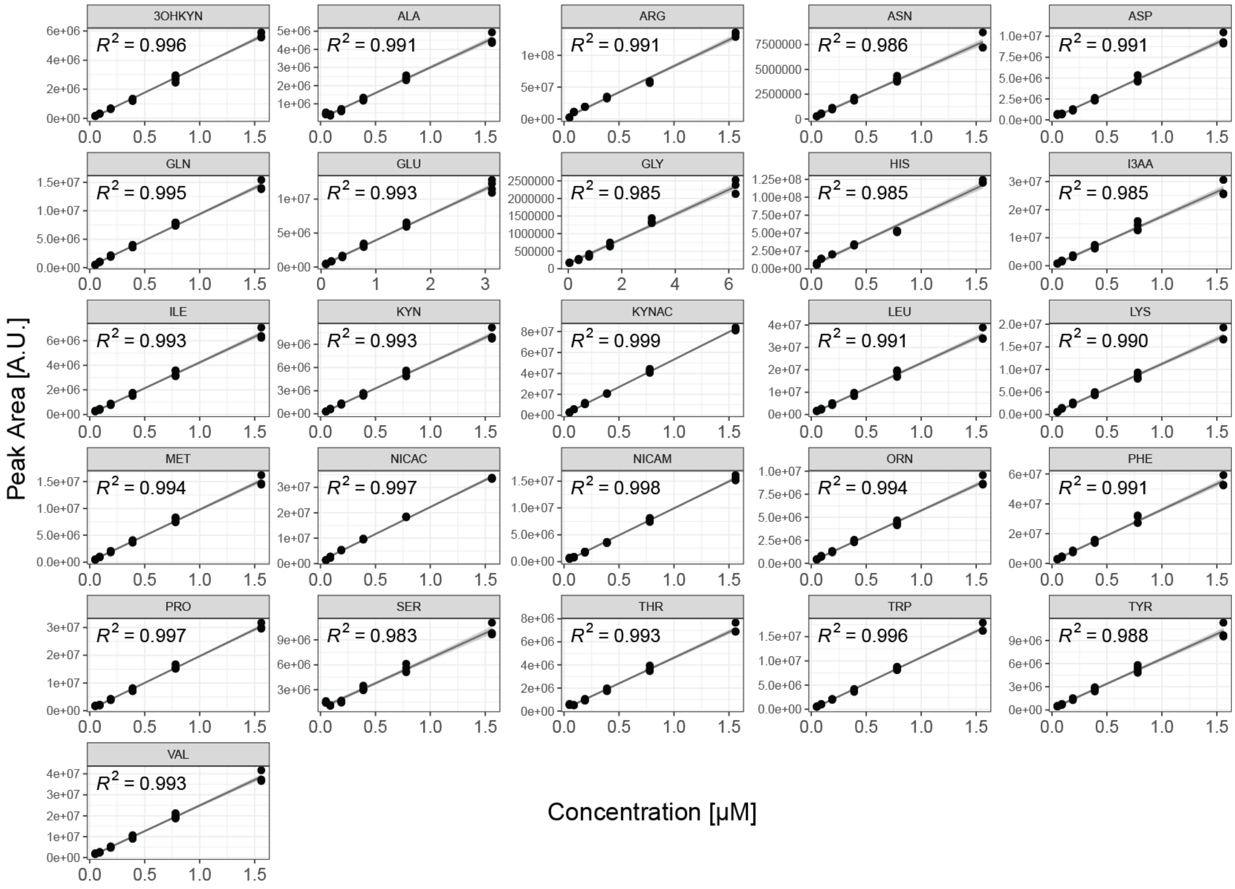


**Figure S3: Coefficient of determination ($R^2$) of the metabolites analyzed using dual-column U(H)PLC-MRM-MS. A dilution series for metabolites from 0.05 µM to and 6.25 µM and linear regression of 12C peak area versus metabolite concentration was used to determine $R^2$ and examine the linearity. n= 4.**

**Table S1.: Stable-isotope labeled canonical and non-canonical amino acids and tryptophan metabolites.**

| Metabolite | Abbreviation | Label, purity | Dissolved in | Stock [mM] | Vendor ID |
|---|---|---|---|---|---|
| 3-Hydroxykynurenine | 3OHKYN | 13C3, 98%; 15N, 98% | 0.1% FA | 15 | CNLM-10399 |
| Alanine | ALY | 13C3, 99%; 15N, 99% | 0.1 N HCl | 2,5 | MSK-CAA-1 |
| Arginine | ARG | 13C6, 99%; 15N4, 99% | 0.1 N HCl | 2,5 | MSK-CAA-1 |
| Asparagine | ASN | 13C4, 99%; 15N2, 99% | 0.1 N HCl | 2,5 | MSK-CAA-1 |
| Aspartate | ASP | 13C4, 99%; 15N, 99% | 0.1 N HCl | 2,5 | MSK-CAA-1 |
| Glutamate | GLN | 13C5, 99%; 15N, 99% | 0.1 N HCl | 2,5 | MSK-CAA-1 |
| Glutamine | GLU | 13C5, 99%; 15N2, 99% | 0.1 N HCl | 2,5 | MSK-CAA-1 |
| Glycine | GLY | 13C2, 99%; 15N, 99% | 0.1 N HCl | 2,5 | MSK-CAA-1 |
| Histidine | HIS | 13C6, 97-99%; 15N3, 97-99% | 0.1 N HCl | 2,5 | MSK-CAA-1 |
| Indole-3-acetic Acid | I3AA | 13C6, 99% | 50% MeOH, 0.1% FA | 30 | CLM-1896 |
| Isoleucine | ILE | 13C6, 99%; 15N, 99% | 0.1 N HCl | 2,5 | MSK-CAA-1 |
| Kynurenic acid | KYNAC | 13C6, 99% | DMSO | 5,1 | CLM-11139 |
| Kynurenine | KYN | 13C10, 99% | 0.1% FA | 15 | CLM-9884 |
| Leucine | LEU | 13C6, 99%; 15N, 99% | 0.1 N HCl | 2,5 | MSK-CAA-1 |
| Lysine | LYS | 13C6, 99%; 15N2, 99% | 0.1 N HCl | 2,5 | MSK-CAA-1 |
| Methionine | MET | 13C5, 99%; 15N, 99% | 0.1 N HCl | 2,5 | MSK-CAA-1 |
| Nicotinamide | NICAC | 13C6, 99% | 0.1% FA | 15 | CLM-9925 |
| Nicotinic Acid | NICAM | 13C6, 99% | 0.1% FA | 15 | CLM-9954 |
| Ornithine | ORN | 13C5, 98% | 0.1% FA | 2,5 | MSK-NCAA-1 |
| Phenylalanine | PHE | 13C9, 99%; 15N, 99% | 0.1 N HCl | 2,5 | MSK-CAA-1 |
| Proline | PRO | 13C5, 99%; 15N, 99% | 0.1 N HCl | 2,5 | MSK-CAA-1 |
| Serine | SER | 13C3, 99%; 15N, 99% | 0.1 N HCl | 2,5 | MSK-CAA-1 |
| Threonine | THR | 13C4, 97-99%; 15N, 97-99% | 0.1 N HCl | 2,5 | MSK-CAA-1 |
| Tryptophan | TRP | 13C11, 99%; 15N2, 99% | 0.1 N HCl | 2,5 | MSK-CAA-1 |
| Tyrosine | TYR | 13C9, 99%; 15N, 99% | 0.1 N HCl | 2,5 | MSK-CAA-1 |
| Valine | VAL | 13C5, 99%; 15N, 99% | 0.1 N HCl | 2,5 | MSK-CAA-1 |

**Table S2: Parameters used for the different extraction workflows to optimize the semi-automated extraction workflow.**

| | **Workflow I** | **Workflow II** | **Workflow III** | **Workflow IV** |
|---|---|---|---|---|
| Sample volume [µL] | 40 | 20 | 40 | 40 |
| Extraction solvent volume [µL] | 160 | 180 | 160 | 160 |
| Incubation on the shaker at 4°C | 5 min, 600 rpm | 5 min, 600 rpm | 5 min, 600 rpm | 5 min, 600 rpm |
| First centrifugation, 4 °C, 4300 x g | 10 min | 10 min | 20 min | 10 min |
| Transfer volume of the supernatant [µL] | 60 | 60 | 60 | 60 |
| Volume of dilution solvent [µL] | 60 | 60 | 60 | 60 |
| Second centrifugation 4 °C, 4300 x g | - | - | - | 10 min |
| Transfer volume of the supernatant [µL] | - | - | - | 60 |
| Incubation on the shaker at 4°C | 5 min, 600 rpm | 5 min, 600 rpm | 5 min, 600 rpm | 5 min, 600 rpm |

**Table S3: Overview of optimized pipetting settings used for semi-automated extraction of amino acids and tryptophan metabolites using the robotic liquid handling platform (Andrew+).**

| | Pipetting Step | | | |
|---|---|---|---|---|
| Pipetting Settings | Transfer of serum to 96 well plate | Transfer of extraction solvent to 96 well plate | Transfer of 0.1% FA to 96 well plate | Transfer of supernatant to 96 well plate |
| Pipetting Mode | Forward | Forward | Forward | Forward |
| Aspiration Speed | Slow | Normal | Normal | Slow |
| Dispensing Speed | Normal | Normal | Normal | Normal |
| Air Cushion | Air bottom cushion (low viscosity aspiration) | Air bottom cushion (low viscosity aspiration) | Air bottom cushion (low viscosity aspiration) | Air bottom cushion (low viscosity aspiration) |
| Pipette moving speed | Normal | Normal | Normal | Normal |
| Tip Position Source | with respect to bottom (avoid touching the bottom) | with respect to bottom (avoid touching the bottom) | with respect to bottom (avoid touching the bottom) | with respect to bottom (avoid touching the bottom) |
| Customized aspiration height | no | no | no | 5mm |
| Tip Position Destination | with respect to liquid | with respect to liquid | with respect to liquid | with respect to liquid |
| Customized dispensing height | no | no | no | no |
| Mixing | no | no | no | no |
| Mechanical arm motion speed | Normal | Normal | Normal | Normal |

**Table S4: Peak widths at half height ($w_{1/2}$) of amino acids and tryptophan metabolites.** Metabolites were separated by mixed-mode chromatography (BEH C18 AX) and reversed-phase chromatography (HSS T3) using 0.1% formic acid (FA) and 0.05% difluoroacetic acid (DFA) as mobile phase modifiers.

| | **BEH C18 AX** | | **HSS T3** | |
|---|---|---|---|---|
| **Metabolite** | **0.1% FA $w_{1/2}$ [sec]** | **0.05% DFA $w_{1/2}$ [sec]** | **0.1% FA $w_{1/2}$ [sec]** | **0.05% DFA $w_{1/2}$ [sec]** |
| 3-hydroxykynurenine (3OHKYN) | 2.34 | 2.30 | 3.35 | 1.45 |
| Alanine (ALA) | 2.62 | 1.89 | 1.58 | 1.84 |
| Arginine (ARG) | 2.26 | 1.44 | 1.99 | 1.96 |
| Asparagine (ASN) | 1.99 | 2.33 | 2.04 | 2.16 |
| Aspartate (ASP) | 2.26 | not detected | 2.02 | 2.04 |
| Glutamine (GLN) | 2.23 | 2.11 | 2.54 | 2.09 |
| Glutamate (GLU) | 2.25 | 2.25 | 2.51 | 2.26 |
| Glycine (GLY) | 1.94 | 1.95 | 1.85 | 2.20 |
| Histidine (HIS) | 2.12 | 1.57 | 2.14 | 2.29 |
| Indole-3-acetic acid (I3AA) | 1.52 | 1.48 | 1.52 | 1.33 |
| Isoleucine (ILE) | 2.13 | 2.29 | 1.96 | 1.72 |
| Kynurenin (KYN) | 1.70 | 1.91 | 2.03 | 1.70 |
| Kynurenic acid (KYNAC) | not detected | not detected | 1.63 | 1.28 |
| Leucine (LEU) | 2.84 | 2.78 | 1.99 | 1.66 |
| Lysine (LYS) | 2.20 | 1.50 | 1.56 | 2.34 |
| Methionine (MET) | 2.08 | 2.28 | 2.26 | 2.93 |
| Nicotinic acid (NICAC) | 2.53 | not detected | 1.55 | 1.97 |
| Nicotinamide (NICAM) | 1.72 | not detected | 3.18 | 2.11 |
| Ornitine (ORN) | 2.19 | 1.41 | 1.55 | 2.20 |
| Phenylalanine (PHE) | 1.31 | 1.54 | 2.11 | 1.34 |
| Proline (PRO) | 2.28 | 2.12 | 2.09 | 1.82 |
| Serine (SER) | 2.14 | 2.17 | 1.78 | 2.20 |
| Threonine (THR) | 1.99 | 2.15 | 2.38 | 2.32 |
| Tryptophan (TRP) | 2.08 | 1.93 | 1.80 | 1.22 |
| Tyrosine (TYR) | 2.21 | 3.31 | 1.89 | 1.67 |
| Valine (VAL) | 2.13 | 2.13 | 2.76 | 2.89 |
| Mean | 2.10 | 2.01 | 2.06 | 1.93 |

**Table S5: Within-run accuracy and within-run precision. Accuracy is represented by the modulus of the mean absolute percentage error (MAPE) and precision by the coefficient of variation (CV) for five concentrations. n= 5.**

| | 0.195 µM | | 0.391 µM | | 3.125 µM | | 4.687 µM | | 6.25 µM | |
|---|---|---|---|---|---|---|---|---|---|---|
| **Metabolite** | **MAPE [%]** | **CV [%]** | **MAPE [%]** | **CV [%]** | **MAPE [%]** | **CV [%]** | **MAPE [%]** | **CV [%]** | **MAPE [%]** | **CV [%]** |
| 3-hydroxykynurenine (3OHKYN) | 4.71 | 4.33 | 2.73 | 1.25 | 2.51 | 2.98 | 2.62 | 2.69 | 3.82 | 1.84 |
| Alanine (ALA) | 4.16 | 4.40 | 3.66 | 4.09 | 1.59 | 1.63 | 2.29 | 2.96 | 3.41 | 3.36 |
| Arginine (ARG) | 5.97 | 3.44 | 4.10 | 4.23 | 3.42 | 2.32 | 1.97 | 2.30 | 4.11 | 4.54 |
| Asparagine (ASN) | 3.91 | 5.34 | 4.17 | 5.04 | 3.96 | 2.32 | 3.80 | 3.96 | 4.35 | 1.74 |
| Aspartate (ASP) | 2.96 | 3.48 | 4.63 | 2.51 | 1.61 | 1.09 | 1.46 | 1.78 | 1.14 | 1.45 |
| Glutamine (GLN) | 7.31 | 7.09 | 9.99 | 10.04 | 10.07 | 12.09 | 7.60 | 9.94 | 8.01 | 8.09 |
| Glutamate (GLU) | 6.55 | 4.88 | 2.94 | 3.35 | 2.99 | 3.72 | 2.73 | 2.96 | 2.21 | 2.99 |
| Glycine (GLY) | 9.28 | 7.36 | 4.75 | 6.29 | 5.43 | 5.06 | 6.78 | 6.65 | 7.18 | 9.59 |
| Histidine (HIS) | 2.98 | 1.18 | 1.02 | 1.18 | 2.65 | 2.26 | 5.40 | 6.21 | 1.65 | 1.80 |
| Indole-3-acetic acid (I3AA) | 2.11 | 2.25 | 3.16 | 2.13 | 2.90 | 1.85 | 2.04 | 2.77 | 1.72 | 2.06 |
| Isoleucine (ILE) | 2.52 | 2.27 | 1.99 | 2.10 | 1.76 | 1.33 | 1.10 | 1.33 | 1.37 | 2.05 |
| Kynurenin (KYN) | 5.50 | 1.99 | 1.35 | 1.61 | 3.43 | 2.05 | 1.49 | 0.99 | 1.58 | 2.07 |
| Kynurenic acid (KYNAC) | 12.35 | 17.24 | 9.21 | 3.70 | 4.69 | 5.52 | 5.36 | 7.60 | 7.42 | 9.40 |
| Leucine (LEU) | 1.35 | 1.44 | 2.21 | 1.70 | 1.26 | 1.60 | 1.17 | 1.70 | 2.83 | 1.53 |
| Lysine (LYS) | 3.05 | 3.82 | 3.71 | 3.30 | 3.76 | 4.45 | 2.35 | 2.68 | 2.24 | 2.73 |
| Methionine (MET) | 4.19 | 2.24 | 2.93 | 2.47 | 3.96 | 3.95 | 1.90 | 2.44 | 1.59 | 2.08 |
| Nicotinic acid (NICAC) | 3.42 | 3.28 | 9.18 | 5.17 | 8.45 | 13.27 | 7.16 | 10.77 | 7.01 | 1.75 |
| Nicotinamide (NICAM) | 20.71 | 5.78 | 3.21 | 2.84 | 6.77 | 9.93 | 9.27 | 11.66 | 12.66 | 12.07 |
| Ornitine (ORN) | 2.97 | 3.13 | 2.03 | 2.29 | 1.30 | 0.88 | 0.91 | 1.12 | 0.81 | 0.91 |
| Phenylalanine (PHE) | 7.06 | 3.59 | 4.85 | 4.27 | 5.09 | 2.19 | 2.09 | 1.77 | 2.04 | 2.49 |
| Proline (PRO) | 4.65 | 3.68 | 5.95 | 2.01 | 1.40 | 1.62 | 1.55 | 1.70 | 2.54 | 2.68 |
| Serine (SER) | 10.91 | 2.94 | 6.56 | 1.04 | 1.76 | 2.28 | 3.63 | 3.81 | 1.61 | 1.67 |
| Threonine (THR) | 3.02 | 4.01 | 2.51 | 2.88 | 1.97 | 1.85 | 1.10 | 1.41 | 3.86 | 2.00 |
| Tryptophan (TRP) | 3.83 | 5.32 | 1.57 | 1.68 | 2.05 | 1.77 | 1.69 | 1.75 | 3.18 | 0.67 |
| Tyrosine (TYR) | 1.29 | 0.91 | 2.40 | 2.35 | 1.38 | 0.94 | 1.08 | 1.43 | 0.97 | 0.94 |
| Valine (VAL) | 4.65 | 5.80 | 1.43 | 2.01 | 1.01 | 1.27 | 1.34 | 1.79 | 2.58 | 3.49 |
| Mean | 5.44 | 4.28 | 3.93 | 3.14 | 3.35 | 3.47 | 3.07 | 3.70 | 3.53 | 3.18 |

**Table S6: Between-run accuracy and between-run precision. Accuracy is represented by the modulus of the mean absolute percentage error (MAPE) and precision is represented by the coefficient of variation (CV) for five concentrations. n= 20.**

| | 0.195 µM | | 0.391 µM | | 3.125 µM | | 4.687 µM | | 6.25 µM | |
|---|---|---|---|---|---|---|---|---|---|---|
| **Metabolite** | **MAPE [%]** | **CV [%]** | **MAPE [%]** | **CV [%]** | **MAPE [%]** | **CV [%]** | **MAPE [%]** | **CV [%]** | **MAPE [%]** | **CV [%]** |
| 3-hydroxykynurenine (3OHKYN) | 3.09 | 3.17 | 2.10 | 2.57 | 1.28 | 1.66 | 2.37 | 2.83 | 2.02 | 2.28 |
| Alanine (ALA) | 9.02 | 5.29 | 4.83 | 4.57 | 3.19 | 2.77 | 1.86 | 2.60 | 3.46 | 2.62 |
| Arginine (ARG) | 5.41 | 4.90 | 3.04 | 3.72 | 2.89 | 2.69 | 2.42 | 2.80 | 2.77 | 3.25 |
| Asparagine (ASN) | 4.23 | 4.99 | 2.27 | 2.87 | 3.12 | 3.28 | 2.44 | 3.19 | 2.63 | 3.72 |
| Aspartate (ASP) | 3.75 | 4.26 | 4.38 | 1.59 | 1.85 | 2.23 | 1.60 | 1.56 | 1.61 | 1.54 |
| Glutamine (GLN) | 5.05 | 5.12 | 3.41 | 4.34 | 1.98 | 2.43 | 2.67 | 3.69 | 3.85 | 3.08 |
| Glutamate (GLU) | 5.36 | 6.61 | 3.12 | 4.03 | 3.00 | 2.97 | 2.07 | 2.30 | 2.30 | 2.75 |
| Glycine (GLY) | 15.49 | 16.82 | 5.31 | 6.30 | 5.41 | 6.72 | 4.45 | 5.21 | 4.95 | 6.25 |
| Histidine (HIS) | 7.56 | 7.75 | 2.45 | 3.13 | 2.88 | 3.23 | 2.71 | 3.30 | 1.53 | 2.06 |
| Indole-3-acetic acid (I3AA) | 3.21 | 4.16 | 1.92 | 2.32 | 3.14 | 2.35 | 2.11 | 2.64 | 2.70 | 2.65 |
| Isoleucine (ILE) | 2.82 | 3.75 | 2.98 | 3.62 | 1.68 | 1.67 | 2.07 | 2.41 | 0.90 | 1.11 |
| Kynurenin (KYN) | 4.31 | 4.28 | 2.82 | 3.41 | 2.49 | 2.74 | 2.48 | 2.92 | 2.29 | 2.69 |
| Kynurenic acid (KYNAC) | 12.56 | 17.96 | 14.96 | 17.72 | 8.65 | 12.19 | 9.59 | 11.58 | 7.64 | 9.02 |
| Leucine (LEU) | 2.61 | 3.16 | 1.60 | 2.00 | 1.69 | 1.92 | 1.23 | 1.43 | 1.44 | 1.75 |
| Lysine (LYS) | 1.87 | 2.05 | 1.75 | 1.63 | 2.61 | 2.39 | 1.39 | 1.57 | 1.62 | 1.85 |
| Methionine (MET) | 3.96 | 4.39 | 2.10 | 2.03 | 2.82 | 3.00 | 2.74 | 3.59 | 2.12 | 2.59 |
| Nicotinic acid (NICAC) | 9.93 | 12.21 | 8.99 | 10.92 | 6.50 | 8.35 | 8.94 | 12.32 | 6.08 | 7.87 |
| Nicotinamide (NICAM) | 16.63 | 20.71 | 14.35 | 15.91 | 11.00 | 15.45 | 12.29 | 21.36 | 4.50 | 6.90 |
| Ornitine (ORN) | 3.65 | 3.17 | 2.86 | 1.62 | 1.54 | 2.12 | 1.32 | 1.78 | 1.04 | 1.28 |
| Phenylalanine (PHE) | 6.87 | 4.58 | 2.06 | 2.70 | 1.93 | 1.64 | 2.11 | 2.67 | 2.70 | 2.46 |
| Proline (PRO) | 3.79 | 4.40 | 2.36 | 3.06 | 2.35 | 3.06 | 2.14 | 2.58 | 2.35 | 2.85 |
| Serine (SER) | 16.68 | 6.76 | 7.32 | 3.54 | 2.72 | 2.81 | 1.62 | 2.00 | 2.16 | 2.49 |
| Threonine (THR) | 5.41 | 4.18 | 3.35 | 3.21 | 2.23 | 2.42 | 1.82 | 2.35 | 3.14 | 1.95 |
| Tryptophan (TRP) | 5.77 | 4.25 | 2.40 | 3.00 | 2.43 | 2.36 | 1.83 | 2.38 | 3.48 | 2.54 |
| Tyrosine (TYR) | 3.97 | 2.43 | 1.57 | 1.95 | 1.90 | 1.90 | 1.79 | 2.15 | 1.56 | 1.59 |
| Valine (VAL) | 3.38 | 3.42 | 2.77 | 3.16 | 3.39 | 3.22 | 2.01 | 2.36 | 2.95 | 3.55 |
| Mean | 6.40 | 6.34 | 4.12 | 4.42 | 3.26 | 3.75 | 3.08 | 4.06 | 2.84 | 3.18 |

**Table S7: Optimization of the semi-automated extraction and sample preparation workflow for the analysis of metabolites in human serum samples.** CV: coefficient of variation (CV). For statistical analysis comparing workflow II and workflow IV, the Mann-Whitney U test was used. n=8 independent experiments.

| | **Normalized metabolites CV [%]** | | **Recovery [%]** | |
|---|---|---|---|---|
| **Metabolite** | **Workflow II** | **Workflow IV** | **Workflow II** | **Workflow IV** |
| Alanine (ALA) | 10 | 12.4 | 72.5 | 88.6 |
| Arginine (ARG) | 11.5 | 9.1 | 71.5 | 84.1 |
| Glutamine (GLN) | 10.7 | 8.6 | 81.7 | 88 |
| Glutamate (GLU) | 17 | 10.1 | 81.7 | 87.6 |
| Glycine (GLY) | 18.7 | 10.2 | 71.6 | 93.2 |
| Histidine (HIS) | 8.4 | 8.8 | 78.3 | 84.3 |
| Isoleucine (ILE) | 8.6 | 7.7 | 84 | 82.7 |
| Leucine (LEU) | 8.2 | 9.1 | 102.9 | 98.9 |
| Lysine (LYS) | 15.1 | 11.7 | 72.3 | 84.4 |
| Methionine (MET) | 8.5 | 6.9 | 79.6 | 89.8 |
| Phenylalanine (PHE) | 3.5 | 3.1 | 89.5 | 95.4 |
| Proline (PRO) | 11.5 | 7.2 | 78.4 | 82.3 |
| Serine (SER) | 9.8 | 10.7 | 79.1 | 89.7 |
| Threonine (THR) | 30.6 | 32.8 | 26.8 | 68.2 |
| Tryptophan (TRP) | 7.8 | 6.9 | 77.4 | 86.7 |
| Tyrosine (TYR) | 9 | 7.7 | 78.5 | 88.7 |
| Valine (VAL) | 8.1 | 8.3 | 78.4 | 84.8 |
| **Mean** | 11.6 | 10.1 | 76.7 | 86.9 |
| **SD** | 6.1 | 6.2 | 14.9 | 6.6 |
| **p-value** | 0.2703 | | 0.0006 | |

**Table S8: Optimization of the semi-automated extraction and sample preparation workflow for the analysis of metabolites in human plasma samples.** CV: coefficient of variation (CV). For statistical analysis comparing the workflow using one clearance step with the workflow using two clearance steps, the Mann-Whitney U test was used. n=8 independent experiments.

| | **Normalized metabolites CV [%]** | | **Recovery [%]** | |
|---|---|---|---|---|
| **Metabolite** | **One clearance step** | **Two clearance steps** | **One clearance step** | **Two clearance steps** |
| Alanine (ALA) | 45.4 | 12.5 | 73.9 | 90 |
| Arginine (ARG) | 62.3 | 8 | 51.7 | 82.7 |
| Asparagine (ASP) | 27.1 | 27.2 | 74.9 | 111.8 |
| Glutamine (GLN) | 229.4 | 6.3 | 90.3 | 92.2 |
| Glutamate (GLU) | 142.2 | 12.9 | 92 | 90.9 |
| Glycine (GLY) | 45.5 | 9 | 78.7 | 89.1 |
| Histidine (HIS) | 56.6 | 7.5 | 62.5 | 96.1 |
| Isoleucine (ILE) | 244.7 | 9.6 | 21.9 | 120.2 |
| Leucine (LEU) | 113.2 | 14.9 | 52.8 | 109.3 |
| Lysine (LYS) | 57.2 | 11.4 | 56.9 | 86.1 |
| Methionine (MET) | 125.8 | 8 | 99.2 | 97.2 |
| Phenylalanine (PHE) | 150.4 | 8.5 | 82.1 | 93.3 |
| Proline (PRO) | 58.8 | 6.6 | 72.6 | 93.4 |
| Serine (SER) | 46.5 | 7.3 | 71 | 90.6 |
| Threonine (THR) | 177.7 | 7.1 | 86.3 | 94 |
| Tryptophan (TRP) | 58 | 15.9 | 76.5 | 92.2 |
| Tyrosine (TYR) | 56.9 | 9 | 26.6 | 112.8 |
| Valine (VAL) | 54.2 | 6.2 | 69.8 | 94 |

| | | | | |
|---|---|---|---|---|
| **Mean** | 97.3 | 10.4 | 68.9 | 96.4 |
| **SD** | 66.6 | 5.1 | 20.8 | 10.2 |
| **p-value** | <0.0001 | | <0.0001 | |

**Table S9: Optimization of the semi-automated extraction and sample processing workflow for the analysis of metabolites in human plasma samples.** CV: coefficient of variation (CV). For statistical analysis comparing the workflow using a 1:10 or a 1:20 sample dilution, Mann-Whitney U test was used. n=8 independent experiments.

| | **Normalized metabolites CV [%]** | | **Recovery [%]** | |
|---|---|---|---|---|
| **Metabolite** | **1:10 dilution** | **1:20 dilution** | **1:10 dilution** | **1:20 dilution** |
| 3-hydroxykynurenine (3OHKYN) | 11.1 | 22.5 | 117.4 | 111.1 |
| Arginine (ARG) | 2.7 | 2.9 | 79.4 | 84.2 |
| Asparagine (ASN) | 2.7 | 3 | 104.7 | 113.4 |
| Aspartate (ASP) | 2.5 | 3.5 | 106.2 | 108.4 |
| Glutamine (GLN) | 3.5 | 3.7 | 109.7 | 117.8 |
| Glutamate (GLU) | 3 | 3.5 | 106.3 | 112.9 |
| Glycine (GLY) | 2.8 | 4.5 | 110.6 | 118.7 |
| Histidine (HIS) | 3.3 | 2.4 | 103.5 | 110.7 |
| Indole-3-acetic acid (I3AA) | 2.7 | 2.8 | 112.3 | 124.2 |
| Isoleucine (ILE) | 2.4 | 2.1 | 108.7 | 119.1 |
| Kynurenin (KYN) | 3.2 | 4.4 | 114.9 | 117.1 |
| Kynurenic acid (KYNAC) | 35.8 | 38.6 | 105.4 | 200.6 |
| Leucine (LEU) | 3 | 2.7 | 109.8 | 117 |
| Lysine (LYS) | 2.5 | 3 | 89.4 | 94.7 |
| Methionine (MET) | 3.1 | 2.4 | 109.9 | 117.6 |
| Phenylalanine (PHE) | 3.6 | 2.8 | 108.9 | 115.8 |
| Proline (PRO) | 2.3 | 3.4 | 107.7 | 116.7 |
| Serine (SER) | 2.2 | 2.6 | 104.5 | 114.9 |
| Threonine (THR) | 1.8 | 4.5 | 104.9 | 114.9 |
| Tryptophan (TRP) | 2.3 | 2.6 | 107.8 | 111.7 |
| Tyrosine (TYR) | 2.4 | 3.3 | 109.9 | 117.7 |
| Valine (VAL) | 3.8 | 2.6 | 107.6 | 116.5 |

| | | | | |
|---|---|---|---|---|
| **Mean** | 4.7 | 5.6 | 106.3 | 117.1 |
| **SD** | 7.2 | 8.5 | 8.0 | 20.5 |
| **p-value** | 0.2085 | | <0.0001 | |

**Table S10: Intra-assay variability of the optimized semi-automated workflow.** Metabolites were extracted from human serum and human plasma. CV: coefficient of variation (CV). n=7 independent experiments. Statistical analysis: Mann-Whitney U test.

| | **Normalized metabolites CV [%]** | |
|---|---|---|
| **Metabolite** | **Serum** | **Plasma** |
| 3-hydroxykynurenine (3OHKYN) | 26 | 8.2 |
| Alanine (ALA) | 6.3 | 4.4 |
| Arginine (ARG) | 21.4 | 4.3 |
| Asparagine (ASN) | 6.6 | 3.4 |
| Aspartate (ASP) | 14.4 | 24.4 |
| Glutamine (GLN) | 8.1 | 4.1 |
| Glutamate (GLU) | 6.4 | 4.9 |
| Glycine (GLY) | 13.3 | 5.2 |
| Histidine (HIS) | 15.1 | 8.3 |
| Indole-3-acetic acid (I3AA) | 6.6 | 4.7 |
| Isoleucine (ILE) | 6.8 | 3.9 |
| Kynurenin (KYN) | 7.4 | 4.8 |
| Kynurenic acid (KYNAC) | 10.9 | 18.2 |
| Leucine (LEU) | 7 | 4.5 |
| Lysine (LYS) | 7.1 | 3.9 |
| Methionine (MET) | 6.6 | 2.9 |
| Nicotinic acid (NICAC) | - | 74 |
| Nicotinamide (NICAM) | 5.5 | 12.7 |
| Ornitine (ORN) | 7 | 4.5 |
| Phenylalanine (PHE) | 6.5 | 2.9 |
| Proline (PRO) | 6.7 | 3.9 |
| Serine (SER) | 7 | 4.2 |
| Threonine (THR) | 6.3 | 3.9 |
| Tryptophan (TRP) | 7 | 3.8 |
| Tyrosine (TYR) | 7.2 | 3.8 |
| Valine (VAL) | 7.9 | 3.7 |

| | | |
|---|---|---|
| **Mean** | 9.2 | 8.8 |
| **SD** | 5.1 | 14.2 |

**Table S11: Inter-assay precision of the optimized semi-automated workflow.** Metabolites were extracted from human serum and human plasma. CV: coefficient of variation (CV). Inter-assay precision was determined for each day and over all 4 days. Statistical analysis: Mann-Whitney U test. n=7 independent experiments per day.

| | **Normalized metabolites human serum CV [%]** | | | | | **Normalized metabolites human plasma CV [%]** | | | | |
|---|---|---|---|---|---|---|---|---|---|---|
| **Metabolite** | **Day1** | **Day2** | **Day3** | **Day4** | **over 4 days** | **Day1** | **Day2** | **Day3** | **Day4** | **over 4 days** |
| 3-hydroxykynurenine (3OHKYN) | 7.32 | 10.41 | 24.34 | 11.88 | 15.36 | 10.18 | - | - | 2.48 | - |
| Alanine (ALA) | 4.56 | 2.58 | 2.7 | 2.72 | 7.28 | 2.46 | 2.32 | 1.11 | 1.27 | 2.04 |
| Arginine (ARG) | 4.81 | 3.51 | 1.64 | 3.4 | 3.8 | 11.47 | 11.47 | 18.76 | 10.29 | 12.21 |
| Asparagine (ASN) | 4.81 | 3.75 | 2.46 | 2.97 | 4.17 | 3.62 | 2.65 | 2.22 | 3.27 | 4.6 |
| Aspartate (ASP) | 9.09 | 3.2 | 4.44 | 3.63 | 5.57 | 2.38 | 3.39 | 2.22 | 5.21 | 3.77 |
| Glutamine (GLN) | 4.09 | 3.29 | 2.95 | 2.57 | 4 | 6.81 | 5.44 | 5.57 | 3.3 | 7.34 |
| Glutamate (GLU) | 9.38 | 4.63 | 9.9 | 8.52 | 12.83 | 4.43 | 8.49 | 7.35 | 7.53 | 8.5 |
| Glycine (GLY) | 4.37 | 3.66 | 4.99 | 34.15 | 20.01 | 0.12 | 7.68 | 6.84 | 3.08 | 8.1 |
| Histidine (HIS) | 3.97 | 4.06 | 2.5 | 3.29 | 4.06 | 3.07 | 3.07 | 6.54 | 7.29 | 4.66 |
| Indole-3-acetic acid (I3AA) | - | - | - | - | - | 5.56 | 4.64 | 5.94 | 5.59 | 8.23 |
| Isoleucine (ILE) | 4.53 | 3.25 | 4.13 | 3.42 | 4.01 | 1.6 | 1.26 | 2.29 | 1.87 | 2.85 |
| Kynurenin (KYN) | 5.26 | 3.38 | 2.71 | 2.3 | 4.26 | 1.05 | 2.55 | 1.15 | 5.53 | 2.71 |
| Leucine (LEU) | 4.53 | 2.54 | 2.03 | 2.53 | 3.64 | 2.87 | 4.44 | 2.29 | 3.1 | 3.3 |
| Lysine (LYS) | 4.84 | 3.75 | 8.12 | 6.23 | 13.67 | 1.49 | 3.62 | 1.4 | 1.73 | 2.4 |
| Methionine (MET) | 3.79 | 3.18 | 3.12 | 2.33 | 3.54 | 1.62 | 1.37 | 1.22 | 2.56 | 3.68 |
| Nicotinic acid (NICAC) | 51.34 | 83.49 | 62.52 | 8.2 | 86.69 | 1.9 | 1.9 | 12.13 | - | - |
| Nicotinamide (NICAM) | - | - | - | - | - | - | - | 18.24 | 4.52 | - |
| Ornitine (ORN) | - | - | - | - | - | 1.85 | 2.18 | 3.9 | 7.46 | 21.88 |
| Phenylalanine (PHE) | 4.19 | 3.28 | 2.91 | 2.43 | 3.65 | 0.68 | 3.98 | 1.77 | 0.81 | 3.37 |
| Proline (PRO) | 4.29 | 2.17 | 4.75 | 3.31 | 4.4 | 4.22 | 3.67 | 1.06 | 1.88 | 3.17 |
| Serine (SER) | 4.43 | 3.24 | 22.03 | 23.3 | 16.84 | 0.49 | 3.16 | 3.5 | 4.37 | 3.56 |
| Threonine (THR) | 2.78 | 4.38 | 5 | 4.26 | 5.03 | 1.38 | 2.88 | 1.97 | 2.54 | 3.73 |
| Tryptophan (TRP) | 4.37 | 3.63 | 3.25 | 3.43 | 3.68 | 3.95 | 4.67 | 4.43 | 1.73 | 3.68 |

| | | | | | | | | | | |
|---|---|---|---|---|---|---|---|---|---|---|
| Tyrosine (TYR) | 4.37 | 2.93 | 4.22 | 3.98 | 3.99 | 1.14 | 0.78 | 0.36 | 4.81 | 3.03 |
| Valine (VAL) | 3.95 | 2.57 | 2.44 | 3.46 | 3.97 | 2.01 | 1.89 | 2.01 | 4.05 | 3.09 |

| | | | | | | | | | | |
|---|---|---|---|---|---|---|---|---|---|---|
| **Mean** | 7.0 | 7.3 | 8.3 | 6.5 | 10.7 | 3.2 | 3.8 | 4.8 | 4.0 | 5.5 |
| **SD** | 10.0 | 17.1 | 13.5 | 7.8 | 17.7 | 2.9 | 2.5 | 5.0 | 2.4 | 4.5 |

**Table S12: Autosampler stability of the metabolites extracted from serum over 72 hours.** The extracts were incubated from 0 to 72 hours (hrs) in the autosampler at 7°C. CV: coefficient of variation [%] for each time point and over 72 hours. n= 3 independent experiments per time point.

| **Metabolite** | **CV [%] metabolites from serum** | | | | **CV [%] over 72 hrs** | **Decrease/ Increase [%]** |
|---|---|---|---|---|---|---|
| | **0 hrs** | **24 hrs** | **48 hrs** | **72 hrs** | | |
| Arginine (ARG) | 0.04 | 2.84 | 2.9 | 4.35 | 3.37 | 0.57 |
| Aspartate (ASP) | 2.17 | 2.25 | 5.55 | 5.47 | 4.31 | 4.93 |
| Glutamine (GLN) | 0.94 | 2.87 | 3.43 | 3.77 | 2.69 | 0.98 |
| Glutamate (GLU) | 0.68 | 4.21 | 4.49 | 4.17 | 3.66 | 2.02 |
| Glycine (GLY) | 0.18 | 13.89 | 4.77 | 17.37 | 11.22 | -5.63 |
| Histidine (HIS) | 0.38 | 1.28 | 1.99 | 5.14 | 2.75 | 0.24 |
| Isoleucine) ILE | 2.83 | 0.84 | 6.15 | 4.85 | 3.83 | 4.38 |
| Leucine (LEU) | 2.83 | 37.35 | 25.74 | 7.32 | 20.42 | 10.59 |
| Lysine (LYS) | 3.2 | 3.04 | 0.77 | 4.94 | 3.52 | 0.50 |
| Methionine (MET) | 1.95 | 2.16 | 5.32 | 5.07 | 3.91 | 4.81 |
| Ornithine (ORN) | 1.63 | 2.63 | 2.91 | 8.61 | 4.79 | -0.07 |
| Phenylalanine (PHE) | 3.38 | 6.33 | 11.97 | 7.37 | 6.51 | -0.05 |
| Proline (PRO) | 3.05 | 6.18 | 4.21 | 4.48 | 4.7 | 2.43 |
| Serine (SER) | 2.87 | 5.52 | 5.01 | 10.75 | 6.51 | -0.58 |
| Threonine (THR) | 1.24 | 5.6 | 4.63 | 9.55 | 6.06 | -2.07 |
| Tryptophan (TRP) | 6.31 | 7.16 | 4.68 | 7.93 | 6.24 | 4.26 |
| Tyrosine (TYR) | 6.33 | 53.61 | 75.34 | 0.1 | 35.7 | 21.01 |
| Valine (VAL) | 1.21 | 1.62 | 4.13 | 4.21 | 3.21 | 3.94 |
| **Mean** | 2.3 | 8.9 | 9.7 | 6.4 | 7.4 | 2.9 |
| **SD** | 1.8 | 13.9 | 17.3 | 3.7 | 8.2 | 5.7 |

**Table S13: Autosampler stability of the metabolites extracted from plasma over 72 hours.** The extracts were incubated from 0 to 72 hours (hrs) in the autosampler at 7°C. CV: coefficient of variation [%] for each time point and over 72 hours. n= 3 independent experiments per time point.

| **Metabolite** | **CV [%] metabolites from serum** | | | | **CV [%] over 72 hrs** | **Decrease/ Increase [%]** |
|---|---|---|---|---|---|---|
| | **0 hrs** | **24 hrs** | **48 hrs** | **72 hrs** | | |
| 3-hydroxykynurenine (3OHKYN) | 0.62 | 1.4 | 2.74 | 13.29 | 6.36 | -4.10 |
| Alanine (ALA) | 1.53 | 1.11 | 2.7 | 1.7 | 2.98 | 0.94 |
| Arginine (ARG) | 4.88 | 19.76 | 4.62 | 10.94 | 10.74 | -5.36 |
| Asparagine (ASN) | 2.09 | 1.89 | 0.04 | 1.4 | 2.62 | 1.63 |
| Aspartate (ASP) | 1.93 | 0.44 | 0.92 | 1.42 | 3.65 | 4.30 |
| Glutamine (GLN) | 1.02 | 1.43 | 0.61 | 2.08 | 2.72 | 0.49 |
| Glutamate (GLU) | 1.18 | 0.92 | 0.12 | 1.35 | 2.46 | 2.64 |
| Glycine (GLY) | 3.12 | 3.14 | 10.07 | 1.13 | 7.06 | -1.26 |
| Histidine (HIS) | 0.67 | 13.98 | 6.46 | 11.22 | 8.75 | -7.82 |
| Indole-3-acetic acid (I3AA) | 1.09 | 0.79 | 1.11 | 2.08 | 2.11 | -0.09 |
| Isoleucine (ILE) | 2.29 | 1.06 | 1.42 | 2.28 | 2.99 | 3.30 |
| Kynurenin (KYN) | 2.04 | 1.29 | 4.31 | 12.9 | 6.44 | -5.16 |
| Leucine (LEU) | 1.56 | 1.31 | 1.22 | 1.83 | 2.94 | 4.07 |
| Lysine (LYS) | 1.37 | 0.15 | 0.84 | 2.08 | 1.58 | -1.73 |
| Methionine (MET) | 1.79 | 1.24 | 0.22 | 1.48 | 3.01 | 4.37 |
| Nicotinic acid (NICAC) | 3.49 | 2.93 | 8.72 | 6.31 | 5.7 | -1.99 |
| Nicotinamide (NICAM) | 2.25 | 3.5 | 0.52 | 7.95 | 4.75 | -0.89 |
| Ornitine (ORN) | 1.37 | 2.73 | 0.02 | 2.43 | 2 | -1.43 |
| Phenylalanine (PHE) | 1.11 | 1.53 | 1.17 | 2.05 | 2.92 | 5.53 |
| Proline (PRO) | 1.65 | 1.86 | 1.12 | 1.91 | 2.29 | 2.73 |
| Serine (SER) | 0.75 | 2 | 0.78 | 0.36 | 2.86 | 3.01 |
| Threonine (THR) | 2.17 | 5.37 | 1.44 | 2.06 | 4.66 | 0.85 |
| Tryptophan (TRP) | 1.77 | 1.82 | 0.54 | 1.11 | 2.65 | 3.70 |
| Tyrosine (TYR) | 0.55 | 0.4 | 1.03 | 2.5 | 2.36 | 3.57 |
| Valine (VAL) | 2.25 | 1.3 | 0.77 | 2.15 | 2.53 | 2.02 |
| **Mean** | 1.8 | 2.9 | 2.1 | 3.8 | 4.0 | 0.5 |
| **SD** | 1.0 | 4.4 | 2.7 | 4.0 | 2.3 | 3.5 |

**Table S14: Quantification of amino acids and tryptophan metabolites extracted from reference plasma sample.** The quantified metabolite levels were compared with the certified values of the NIST SRM 1950 reference sample (*). Mean absolute error (MAE) was calculated as the average of the subtraction of each individual calculated concentration (n= 7 independent experiments) from the provided reference value/certified value. Mean absolute percentage error (MAPE) was calculated as the average of the subtraction of the reference value/certified value from each individual calculated concentration (n=7 independent experiments) divided by the calculated concentration as described by Gray et al [1]. Official Certificate of Analysis is available at www.nist.gov.

| Compound | NIST SRM 1950 Reference Values, (*) Certified Values [µM] | Absolute Quantification median ± SD [µM] | Absolute Quantification mean ± SD [µM] | CV [%] | MAE [µM] | MAPE [%] |
|---|---|---|---|---|---|---|
| Alanine (ALA) | 300 ± 26 | 259.19 ± 7.58 | 258.3 ± 7.58 | 2.93 | 41.7 | 16.2 |
| Arginine (ARG) | 81.4 ± 2.3 (*) | 98.09 ± 8.43 | 101.72 ± 8.43 | 8.29 | 20.3 | 19.6 |
| Asparagine (ASN) | - | 33.58 ± 2.1 | 33.4 ± 2.1 | 6.29 | - | - |
| Aspartate (ASP) | - | 12.87 ± 2.9 | 13.6 ± 2.9 | 21.35 | - | - |
| Glutamine (GLN) | - | 401.79 ± 12.72 | 407.01 ± 12.72 | 3.13 | - | - |
| Glutamate (GLU) | 67.4 ± 18 (*) | 103.28 ± 8.34 | 100.62 ± 8.34 | 8.29 | 33.2 | 32.6 |
| Glycine (GLY) | 245 ± 16 | 263.92 ± 29.28 | 277.18 ± 29.28 | 10.56 | 32.2 | 10.8 |
| Histidine (HIS) | 72.6 ± 3.6 | 74.96 ± 3.9 | 73.95 ± 3.9 | 5.27 | 1.4 | 1.6 |
| Indole-3-acetic acid (I3AA) | | 1.01 ± 0.06 | 1.03 ± 0.06 | 5.41 | - | - |
| Isoleucine (ILE) | 55.5 ± 3.4 | 61.89 ± 2.7 | 61.71 ± 2.7 | 4.38 | 6.2 | 9.9 |
| Kynurenin (KYN) | - | 0.93 ± 0.07 | 0.95 ± 0.07 | 6.99 | - | - |
| Kynurenic acid (KYNAC) | - | 0.05 ± 0.01 | 0.05 ± 0.01 | 11.37 | - | - |
| Leucine (LEU) | 100 ± 6 | 111.91 ± 3.64 | 112.93 ± 3.64 | 3.22 | 12.5 | 11.0 |
| Lysine (LYS) | 140 ± 14 | 139.47 ± 5.06 | 138.75 ± 5.06 | 3.65 | 1.3 | 1.0 |
| Methionine (MET) | 22.3 ± 1.8 | 22.89 ± 0.74 | 22.91 ± 0.74 | 3.22 | 0.6 | 2.6 |
| Ornithine (ORN) | 52.1 ± 2.8 (*) | 57.96 ± 2.5 | 58.56 ± 2.5 | 4.27 | 6.5 | 10.9 |
| Phenylalanine (PHE) | 50.8 ± 7 (*) | 58.62 ± 2.8 | 59.35 ± 2.8 | 4.72 | 8.4 | 13.9 |
| Proline (PRO) | 177 ± 9 | 189.71 ± 5.54 | 189.78 ± 5.54 | 2.92 | 12.8 | 6.7 |
| Serine (SER) | 95.9 ± 4.3 | 103.13 ± 5.33 | 104.89 ± 5.33 | 5.08 | 9.0 | 8.4 |
| Threonine (THR) | 119 ± 6 | 118.49 ± 4.64 | 119.09 ± 4.64 | 3.89 | 0.4 | 0.5 |

| | | | | | | |
|---|---|---|---|---|---|---|
| Tryptophan (TRP) | - | 41.75 ± 1.45 | 42.03 ± 1.45 | 3.44 | - | - |
| Tyrosine (TYR) | 57.3 ± 3 | 58.6 ± 1.96 | 58.17 ± 1.96 | 3.37 | 0.9 | 1.4 |
| Valine (VAL) | 182 ± 10 | 179.54 ± 4.55 | 180.64 ± 4.55 | 2.52 | 1.6 | 0.9 |

1 Gray, N. et al. High-Speed Quantitative UPLC-MS Analysis of Multiple Amines in Human Plasma and Serum via Precolumn Derivatization with 6-Aminoquinolyl-N-hydroxysuccinimidyl Carbamate: Application to Acetaminophen-Induced Liver Failure. Anal Chem 89, 2478-2487, doi:10.1021/acs.analchem.6b04623 (2017).

**Table S15: Analysis of variances (ANOVA) for prostate cancer study.** p-values per metabolite determined by ANOVA including all patient conditions and controls.

| **Metabolite** | **p-value** |
|---|---|
| Alanine (ALA) | 0.1020 |
| Arginine (ARG) | 0.2297 |
| Asparagine (ASN) | 0.1887 |
| Aspartate (ASP) | 0.3752 |
| Glutamine (GLN) | 0.9975 |
| Glutamate (GLU) | 0.8591 |
| Glycine (GLY) | 0.3847 |
| Histidine (HIS) | 0.8924 |
| Isoleucine (ILE) | 0.3837 |
| Leucine (LEU) | 0.1868 |
| Lysine (LYS) | 0.1825 |
| Methionine (MET) | 0.0489 |
| Ornithine (ORN) | 0.0260 |
| Phenylalanine (PHE) | 0.0827 |
| Proline (PRO) | 0.5386 |
| Serine (SER) | 0.1226 |
| Threonine (THR) | 0.8198 |
| Tryptophan (TRP) | 0.1831 |
| Tyrosine (TYR) | 0.0223 |
| Valine (VAL) | 0.3907 |
